# “Let’s Be Black Excellence”: How Black Students Navigate Exclusionary and Affirming Racialized Peer Interactions in Active Learning

**DOI:** 10.64898/2026.06.21.733226

**Authors:** Hannah Nichols, Talia Swanson, Jordan Roberts, Renette-Kaire Tchocksi, Matthew Turnipseed, Chase Anderson, Tatiane Russo-Tait

## Abstract

Racial equity remains a critical challenge in postsecondary science education, as Black students experience higher attrition rates and diminished well-being compared to their White peers. While active learning has been shown to reduce failure rates and narrow achievement gaps, the interpersonal affordances and constraints of peer discussions within these settings remain underexplored for Black students. Grounded in Critical Race Theory and utilizing the analytical lenses of racial microaggressions, microaffirmations, and Community Cultural Wealth, this study addresses this issue by investigating the racialized interpersonal experiences of Black students in active learning college science classrooms. Through semi-structured interviews, this study explores the nature of both exclusionary and affirming peer interactions and how students navigate these dynamics. Findings reveal that Black students frequently encounter racialized microaggressions—manifestations of macro-level anti-Blackness— in the classroom which contribute to isolation and racial battle fatigue. Conversely, students describe instances of microaffirmations, predominantly through small counterspaces created by and for other students of color, which validate their intellectual contributions and foster a sense of belonging. Despite facing these tensions, participants advocate for active learning as a beneficial practice, provided that instructors implement explicit, equitable structures and facilitate culturally responsive classroom climates. These findings offer actionable implications for researchers and practitioners to design more inclusive active learning environments by explicitly addressing interpersonal dynamics, promoting cultural competence, and co-constructing humanizing science classroom climates.

## INTRODUCTION

Racial equity in postsecondary science remains an issue with Black students leaving STEM majors or dropping out of college altogether at higher rates than White students (Riegle-Crumb et al., 2019). For Black students who succeed, they do so at a cost to their health and well-being (McGee, 2016; 2020; 2021). Traditional teaching practices such as solely lecturing contributes to this attrition (Theobald et al., 2020). To reform traditional pedagogies, active learning strategies were introduced and have been shown to reduce failure rates by 55% (Freeman et al., 2014). Active learning can also narrow the achievement gap between marginalized and overrepresented students (Theobald et al., 2020; Haak et al., 2011). Yet, research has also begun to identify how a major component of active learning, peer discussion, has interpersonal affordances and constraints. For example, students such as women, LGBTQIA+ students, students with disabilities, and Christian students report negative experiences related to their marginalized identities such as fear of negative evaluation and lower participation (Reinholz et al., 2020; Cooper & Brownell, 2016; Gin et al., 2020). These students also report positive experiences such as being able to correct misconceptions about their identities or meeting others with similar identities. While this body of work has yet to focus on the racialized experiences of Black students during peer discussions in college science classrooms, it supports the argument for the need to explore the specific affordances and constraints for these students in active learning. This is important because scholarship on the experiences of Black students in STEM higher education broadly, indicates that racialized exclusion and discrimination are also likely happening in peer discussions specifically, leading to inequitable experiences for these students. Therefore, there is a need to explore the positive and negative racialized experiences of Black students in peer discussion to understand what these experiences are and how they can be improved.

## LITERATURE REVIEW

Active learning is an umbrella term for teaching strategies that help students build on their prior knowledge and apply higher order thinking to solve problems, individually and with peers, while also becoming more reflective about their own learning (Lavi et al., 2024). This student-centered approach has been empirically shown to improve knowledge retention, student engagement, interactivity between students, and grades (Prince, 2004; Allsop et al., 2020). In fact, using active learning practices instead of solely lecturing in STEM courses has been shown to decrease student failure rates by 55% when compared to only lecture (Freeman, et al. 2014). Further, students of color and students from low-income backgrounds benefit from active learning more than their overrepresented counterparts, therefore narrowing the equity gaps for marginalized students (Theobald et al., 2020; Haak et al., 2011). Given this research, scholars have suggested that using traditional teaching practices is tantamount to discrimination and that using active learning practices instead would help diversify STEM (Handelsman, et al., 2022).

Recently, however, questions about equity in active learning have arisen. For example, although active learning can shrink equity gaps between students of color and White students, there are cases where it has exacerbated these achievement gaps instead (Theobald et al., 2020). This could be because White students may feel a sense of belonging in their science courses compared to students of color (Theobald et al., 2020). However, because active learning encompasses a myriad of curricular and instructional practices, more studies are needed to understand what specific components of the pedagogy can positively or negatively affect the outcomes of students from racially minoritized backgrounds (Freeman et al., 2014; Theobald et al., 2020). One such line of inquiry has been to explore how marginalized and minoritized identities may become salient and affect the experiences of students from those backgrounds during the peer discussion portion of active learning (e.g. Cooper and Brownell, 2016; Maries et al., 2020; Gin et al., 2020; Edwards et al., 2024; Pfeifer et al., 2024).

Marginalized identities can become salient for students during peer interactions for a variety of reasons, including societal stereotypes and some of the problematic aspects of the culture of science. Western science was historically designed for and by white able-bodied cishetero men from elite backgrounds. Therefore, the culture of science often centers the values, norms, and practices from members of those social positions as the standard to which all others must be compared (Author, 2026; NASEM, 2016). This culture can be exclusionary for students who are not from those backgrounds and affect them in their academic and social experiences, which can lead to attrition from STEM majors (NASEM, 2016; 2023). As college science courses are increasingly adopting active learning strategies that center peer discussions during lecture (e.g., “think-pair-shares” and group work), and given the hypothesis that marginalized identities can become salient during peer interactions, discipline-based education research (DBER) scholars have documented the affordances and constraints of peer discussions for marginalized students in college science classrooms over the past decade.

The extant scholarship shows how peer discussions in science classrooms can be exclusionary for women, LGBTQAI+ students, and students with disabilities to name a few. For example, because women are stereotyped as intellectually inferior compared to men, this can trigger stereotype threat. Stereotype threat is a fear (either conscious or unconscious) of conforming to a negative stereotype, leading to underperformance in evaluative situations (Steele, 1997). It can occur in peer discussion, leading to lower participation rates in male dominated groups, which is linked to lower performance (Aguillon et al., 2020). In addition, LGBTQIA+ students report sensing unspoken homophobic messages likely due to the cisheteronormative culture of science. This increases identity salience in peer discussions, making students worry about coming out and risking avoidance from peers; concerns which can affect cognitive load (Cooper & Brownell, 2016). Further, students with disabilities are nervous about working in groups and having to share their disability status with peers and risk being deemed incompetent (Pfeifer et al., 2023; Gin et al., 2020). Latiné students reported racial microaggressions and classroom isolation (Leyva, et al., 2026). Finally, although being Christian is not a marginalized identity in the US, students’ religious identity can become salient in the science classroom. Christian students report that peer discussions increase a fear of being deemed incapable of doing science and being associated with actions of their church (Edwards et al., 2024). Therefore, due to different aspects of the supposed identity neutral culture of science (Author, 2026; Leyva et al., 2022), peer discussions can increase the salience of a marginalized identity and lead to negative experiences in active learning.

Despite the exclusionary culture of science, research has also highlighted positive experiences of marginalized students in peer discussion. For example, when peer discussion is implemented with powerful community standards or when women work in groups, they report better experiences, and research shows higher performance outcomes (Maries et al., 2020). Students who identify as LGBTQIA+ share that peer discussions provide the opportunity for meeting other members of their community as well as educating non-LGBTQIA+ peers about their identity and shifting their mindsets (Cooper & Brownell, 2016). Furthermore, students with disabilities report changing their motivation to learn after working in groups and valuing being able to ask peers about difficult topics (Gin et al., 2020; Pfeifer et al., 2023). Latiné students experienced racially affirming community building that validated their experiences (Leyva et al., 2026). Christian students report finding community with fellow Christian students, help peers feel comfortable sharing their own religious identities, and can correct misconceptions peers may have about their religion (Edwards et al., 2024). Therefore, peer interactions in active learning science classrooms can also provide a space for positive experiences.

While scholarship has reported on the experiences and perspectives of Black students in active learning as part of broader projects (Cedillo, 2018; Eddy et al, 2017; Eddy & Hogan, 2014; Packard & Sand, 2024; Dancy et al., 2020; Dortch & Patel, 2017; Priddie, 2024; Stanton et al., 2022) to our knowledge, there has yet to be a study intentionally examining Black science students affirming and exclusionary racialized experiences during peer interactions. Yet, this body of work, as well as copious evidence from other types of research on the Black experience in higher education supports the argument for the need to explore the affordances and constraints for Black students during peer interactions. For example, Black students must contend with microaggressions, stereotyping, and discrimination while also pursuing their academic goals (NASEM, 2023). Most institutional norms, values, and practices of higher education in the U.S. are deeply rooted in anti-Blackness. Anti-Blackness is more than racism against Black people, it transcends lack of resources and material inequities and refers to how the American educational system pathologizes Black students as inherent threats and position Blackness as antithetical to the “ideal student” (Dumas & Ross, 2016). Cedillo (2018) further argues that “progressive” inquiry-based practices in STEM, operate as anti-Black constructs by reinforcing a Eurocentric epistemology that delegitimize Black students as people who can be science knowers or knowledge producers. Thus, anti-Blackness is embedded in the culture of science, where several values, norms, and practices subject Black students to hostile and inequitable learning environments (Author, 2026; Diamond & Gomez, 2023; Dumas & Ross, 2016; Castro, 2014; Morton et al., 2023; NASEM, 2023). Black students contend with stereotypes of intellectual and cultural inferiority, with several other discriminatory barriers that exclude them or force them to assimilate to fit White cultural norms (Diamond & Gomez, 2023; Morton et al., 2023).

Yet, Black students are persistent as extant studies also show that they resist racial bias in STEM by finding and creating counterspaces where they engage in racialized interactions that affirm, validate, and build community (Ong et al., 2017; Rolón-Dow & Davison, 2021). Many maintain high achievement in courses and use stereotype management to navigate racialized interactions. Stereotype management is a deliberate effort to track, deflect, or actively disprove the racialized assumptions people hold about them (McGee & Martin, 2011; McGee, 2016; 2021). For example, Black students often engage in code-switching, which involves changing speaking style or appearance to better fit in the White cultural norms of the environment (Woolard et al., 2004; Molinsky, 2007). Some Black science students see code switching as a survival mechanism to navigate White spaces, while others say it is an asset that allows them to easily communicate (Stanton et al., 2022). Regardless of how students interpret these experiences, research shows that these practices can ultimately impact mental and physical health and lead to racial battle fatigue (McGee & Martin, 2011; McGee 2016, 2020; Smith et al., 2007). Racial battle fatigue is the body’s mental and physiological response to chronic stress navigating racial bias, discrimination, and microaggressions over time (McGee, 2020; Smith et al., 2007). It includes affective reactions such as anger, frustration, feat, and helplessness as well as physical symptoms such as increased heart rate and blood pressure, anxiety and depression, to name a few (McGee, 2021; Smith et al., 2007).

The evidence described so far shows that peer discussion in active learning college classrooms can be a site of exclusion and affirmation for students from various marginalized backgrounds such as women, LGBTQIA+ students, religious students, and students with disabilities. While there is evidence that Black students experience discrimination and exclusion, as well as affirmations, in higher education and STEM, these racialized experiences have not yet been systematically explored in the context of peer interactions in active learning college science classrooms. This study seeks to answer the following research questions:

1. What are the exclusionary racialized experiences of Black students during peer discussions in active learning college science classrooms?
2. What are the affirming racialized experiences of Black students during peer discussions in active learning college science classrooms?
3. How do they navigate these affirming and exclusionary experiences in the classroom?
4. Given those experiences, do they think active learning is beneficial? Do they prefer lecture-based courses?
5. What types of instructional practices and structures do they think would facilitate the design of more equitable experiences in peer discussion?

By answering these questions, we hope to elevate the voices of Black students and highlight their experiences–– both positive and negative–– so that we can identify the tensions and possibilities of peer interactions in active learning, which will, in turn, inform the design of racially equitable active learning STEM college science classrooms.

## THEORETICAL FRAMEWORK

### Critical Race Theory

To properly ground this study and support its rationale and research design, we framed it using Critical Race Theory (CRT). CRT has several tenets, and for the purposes of this study, we focus on six: 1) racism is normalized and permanent in the U.S.; 2) the interdisciplinary perspective; 3) racism intersects with other forms of subordination; 4) the centrality of experiential knowledge; 5) the challenge to dominant ideology; and 6) commitment to social justice.

CRT positions racism as endemic and permanent in the lives of Americans, meaning it is in our institutions, systems, laws, and here to stay unless exposed (Yosso et al., 2009). It also necessarily centers an interdisciplinary perspective. Scholars from across disciplines who explore race and racism through sociopolitical and historical lenses argue that racism works as a social system that allocates different forms of power to White people which shape outcomes for marginalized individuals (Bonilla-Silva, 2021). Therefore, arguing that racism is embedded in U.S. society and because of this, it appears through everyday practices including in educational systems (Diamond & Gomez, 2023). In order to better understand how this system manifest in society and in our lives, it is helpful to draw from interdisciplinary research and imagine it occurring at different levels: the macro-level (e.g., policies, culture, ideologies, and stereotypes), meso-level (e.g., institutions and organizations) and micro-level (i.e., intrapersonal and interpersonal). As mentioned, since racism operates at higher education institutions (i.e., the meso-level), it can be inferred that it also manifests at the micro-level in classrooms, particularly during interpersonal interactions that occur via peer discussions. A CRT framing allowed us to predict that students of color are likely to experience racist behavior and harm in the classroom. Further, it compelled us to draw from scholarship outside of biology education research to enrich our understandings of the multiple way racism manifests across systems, including the science pathway. Science culture is known to be particularly exclusionary for people of color while due to its white supremacist and colonial roots (Morton et al., 2023). As our participants major in the sciences, this is an even more pertinent reason to consider the endemic nature of racism for this study.

CRT allowed us to better understand the experiences of students of color at the intersection of other subordinated social identities. Intersectionality encourages researchers to consider the relation of multiple identities (e.g. gender, race, class, disability, religion) and how they lead to inequality and discrimination (Crenshaw, 2013). While intersectionality was added to the tenets of CRT a bit later, it is a vital concept that reveals how single axis thinking undermines struggles for social justice (Gillborn, 2015; Cho et al., 2013). The double-bind is a particularly relevant example of how intersectional oppression can be bigger than racism and sexism alone in the context of STEM (Malcolm et al., 1976). While we did account for the intersectional experiences of our students, racial and gender identities along with first generation status were the most discussed among experiences. Using the framework of CRT allowed for us to critically analyze the experiences of the students along with all social identities that they possess and that participants feel play a role in how they experience peer discussion. CRT centers experiential knowledge and challenges dominant ideology through counterstorytelling and pushing back against color-evasive ideology, respectively. Counterstorytelling allowed this study to be told through the testimonies of students of color and challenge the race-neutral approaches to implementing peer discussion in active learning. Participants shared their own stories of interpersonal experiences of affirmation and exclusion in peer discussion. Therefore, CRT provided further rationale to center students’ voices through interviewing, while simultaneously challenging the dominant narrative that something might be wrong with them, as opposed to the active learning environment (Delgado, 1989; Solorzano & Yosso, 2002).

Finally, by situating our study on a framework that defines race/racism as a social structure with multiple levels, we could examine how micro-level exclusionary or inclusionary interactions in peer discussion can affect students in ways that challenge or reproduce racial inequities and importantly, maintain CRT’s commitment to social justice. By examining the classroom structures at the meso-level, we identified potential levers for change in active learning environments that can make these courses more equitable for students. We also conducted this study foregrounding macro-level issues such as White supremacy, stereotypes, color-evasion, and the values and norms of STEM, lead to potential constraints for peer discussions. The ultimate goal of this study being to contribute to scholarship that will lead to a more socially just design of active learning classrooms.

### Analytical Lenses

#### Microaggressions

Research shows that interpersonal racism rarely occurs in overt forms in contemporary society and instead tends to manifest in interpersonal interactions through racial microaggressions. Grounded in CRT, Solórzano and Pérez Huber (2020) define racial microaggressions as a form of systemic everyday verbal or nonverbal assaults directed towards people of color based on race and such assaults are cumulative and take a psychological, physiological, and academic toll on people of color (p. 6-7). Students of color face microaggressions from peers in universities in a number of ways: racial segregation in informal study groups, classroom isolation, assumptions about intelligence and ability, the devaluing of contributions and ideas, racial jokes, gaslighting, and nonverbal aggressions such as avoidance (Solórzano et al., 2000; Yosso, 2009). It is important to note that microaggressions are not called “micro” because their harm is small. They are called ’micro’ because they happen at the micro-level—meaning person-to-person, everyday interactions. They are the interpersonal manifestations of macro-level/societal systemic oppression (i.e., White supremacy, anti-Blackness). Because our study is investigating the racialized micro-level classroom interactions, it made sense to examine Black student experiences through the analytical lenses of microaggressions.

#### Microaffirmations

Studies show that Black students resist racial microaggressions via racial microaffirmations. Also grounded in CRT, racial microaffirmations are behaviors, verbal remarks, and environmental cues that affirm students of color, acknowledge racialized realities, resist racism, or advance racial justice (Rolón-Dow & Davison, 2021; Solórzano & Pérez Huber 2020). Racial microaffirmations can protect, validate, and recognize the experiences of students of color, helping them feel a sense of belonging and community. Microaffirmations help student integration into the science community (Estrada, 2019). They also help students of color to feel protected and seen when their experiences with racism are acknowledged or when others stand up for them (Rolón-Dow & Davison, 2021). Therefore, we used microaggressions as an analytic lens to explore positive and affirming racialized student experiences.

#### Community Cultural Wealth

We utilize Community Cultural Wealth (CCW) as the final analytical lens for this study. Community cultural wealth is also derived from CRT and highlights the tools and skills or “cultural capitals” that people of color utilize to resist and persist in the face of oppression (Yosso, 2005). Yosso identified six cultural capitals—aspirational, familial, resistant, social, linguistic, and navigational—collectively representing the inherent community assets, communicative skills, and kinship networks that people of color leverage to endure and succeed, despite systemic and structural injustice. Given the pervasiveness of anti-Blackness, we believe it is essential to use CCW to highlight the multitude of assets and strengths Black students use to navigate their positive and negative interpersonal experiences in peer discussion.

By examining peer interactions through these analytical lenses grounded in critical race theoretical framework, we were able to contextualize micro-level (interpersonal) experiences in the meso-level (STEM structure, culture, and climate) as informed by the macro-level (e.g., white supremacy, anti-Blackness).

## METHODOLOGY

Because this study is grounded in Critical Race Theory, it adopts a critical race epistemology (Delgado & Bernal, 2002). This framework rejects the illusion of value-free research, instead acknowledging that knowledge production is inherently political. By operating from this epistemological stance, our study intentionally privileges the experiential knowledge and counter-narratives of Black participants as foundational truths necessary to disrupt dominant, anti-Black discourses in STEM.

### Study Context

The study was conducted in a Research Intensive, Southern predominantly white institution (PWI) in a state with population of 58.7% White, 33.2% Black, 4.9% Asian, and 11.1% Latino residents (U.S. Census Bureau, 2023). Yet, the university’s student population is comprised of 65.1% White students, 10.4% Asian students, 7.57% Black or African American students, and 6.66% Hispanic or Latino students (Data USA, 2022). Most students attending the school are in-state residents, migrating to the town from various high schools––most of which are highly segregated, structurally or geographically. Because of this segregation at the K-12 level, many students of color and White students have never had the opportunity to meaningfully interact with each other until they get to this university. Students of color can experience a culture shock when transitioning to a PWI like this one, while White students might not know how to properly engage with students of color, and their actions might be informed by stereotypes. Another important consideration for this research site is that the institution is considerably invested in institutionalizing active learning by increasing the number of active learning classrooms and institutionalizing professional development in active learning for instructors across the institution. Therefore, recruiting students with experiences in active learning science classrooms will not be difficult, and given the sociohistorical context of the state, we hypothesized that racialized experiences would also be commonplace.

### Participants and Sampling

This study was approved and declared exempt by the institution’s IRB Review [00006761]. Inclusion criteria for this study included being a science major and having completed at least 3 active learning courses that utilize peer discussion at the institution. We completed a total of 58 interviews which averaged about one hour. For the present study we focused on the interviews conducted with self-identified Black and African American students, N=21; 3 Black men and 18 Black women (see rationale for this decision under Data Analysis below). Note that while we asked a series of demographic information from all participants (gender, race, ethnicity, disability status, generation to college status, LGBTQIA+ status), we do not list all intersectional identities on the table and instead only list those identities that are salient for each specific participant to protect their anonymity (i.e., since there are so few Black students in the institution and STEM majors, if we list *all* their identities and their major, we risk identifying the participant). Table 1. Study Participants

**Table 1.**
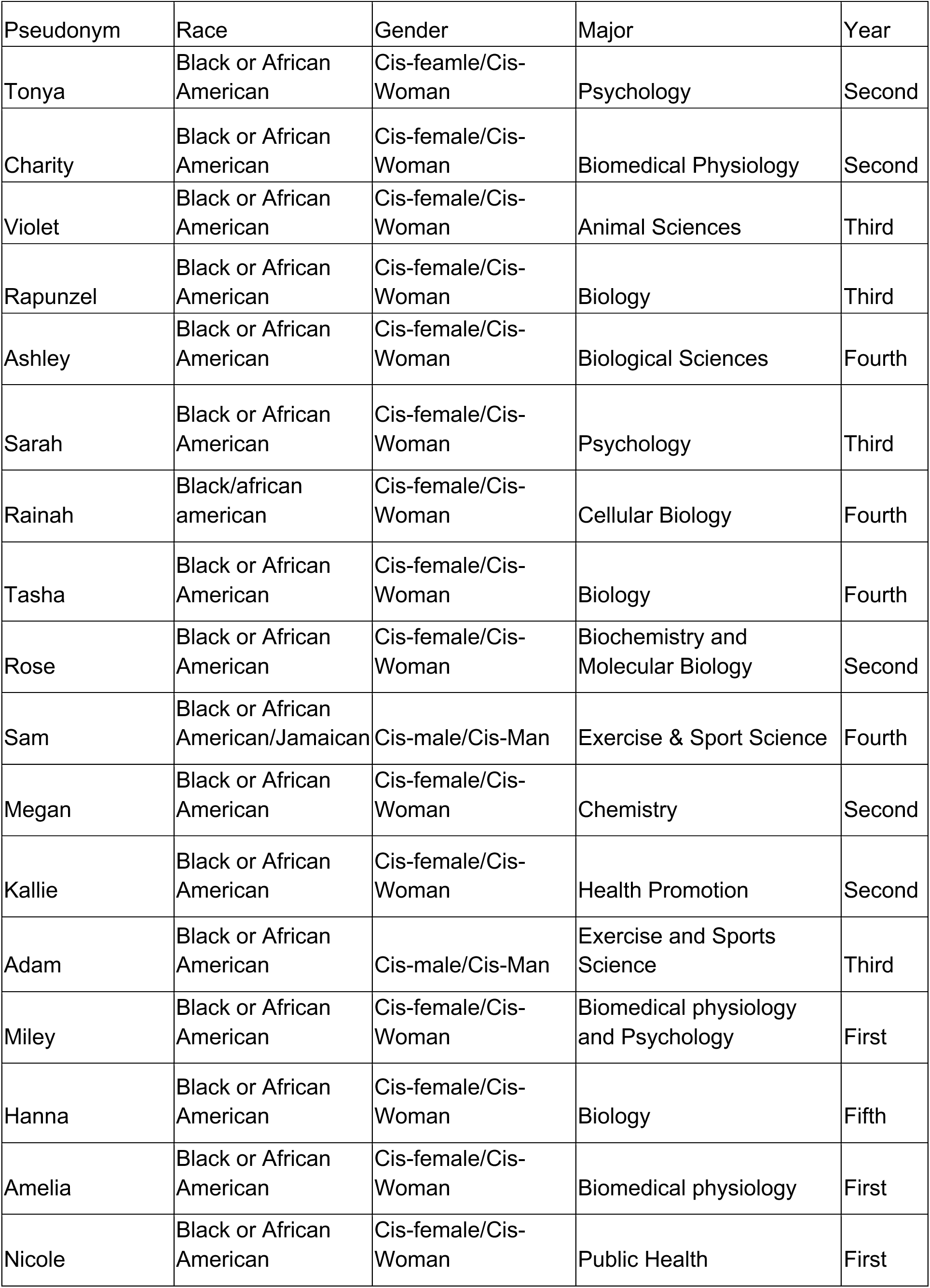
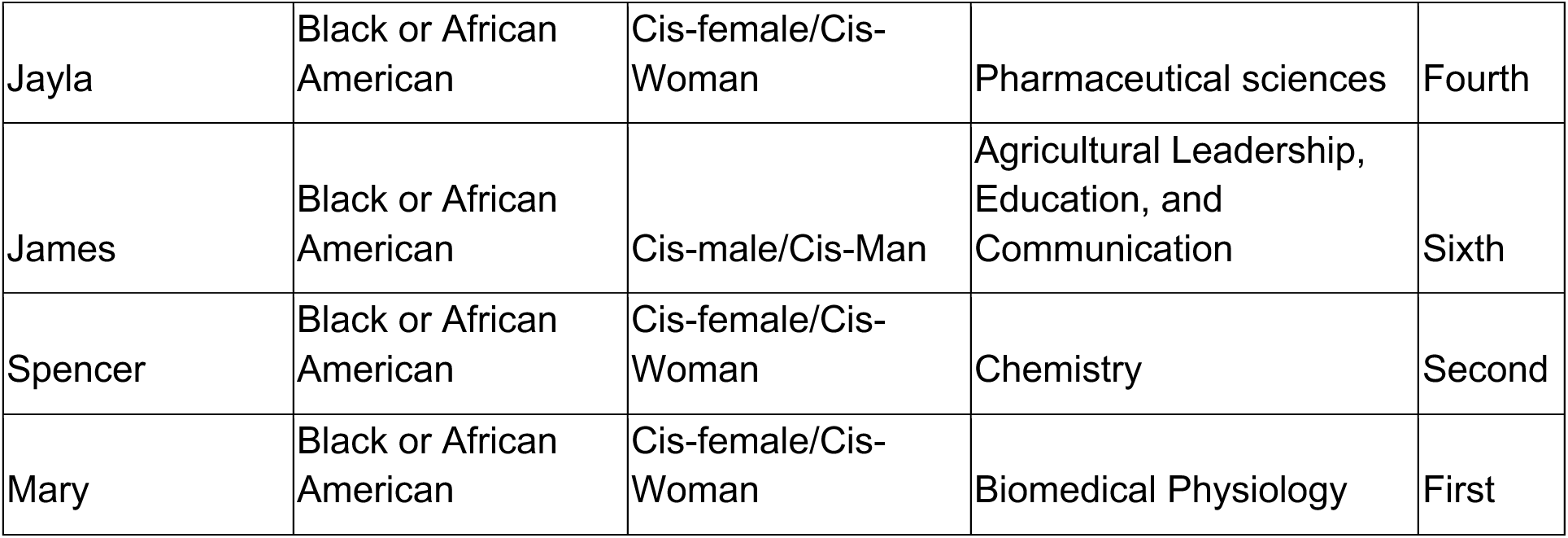
Participant information.

| Pseudonym | Race | Gender | Major | Year |
| --- | --- | --- | --- | --- |
| Tonya | Black or African American | Cis-female/Cis-Woman | Psychology | Second |
| Charity | Black or African American | Cis-female/Cis-Woman | Biomedical Physiology | Second |
| Violet | Black or African American | Cis-female/Cis-Woman | Animal Sciences | Third |
| Rapunzel | Black or African American | Cis-female/Cis-Woman | Biology | Third |
| Ashley | Black or African American | Cis-female/Cis-Woman | Biological Sciences | Fourth |
| Sarah | Black or African American | Cis-female/Cis-Woman | Psychology | Third |
| Rainah | Black/african american | Cis-female/Cis-Woman | Cellular Biology | Fourth |
| Tasha | Black or African American | Cis-female/Cis-Woman | Biology | Fourth |
| Rose | Black or African American | Cis-female/Cis-Woman | Biochemistry and Molecular Biology | Second |
| Sam | Black or African American/Jamaican | Cis-male/Cis-Man | Exercise & Sport Science | Fourth |
| Megan | Black or African American | Cis-female/Cis-Woman | Chemistry | Second |
| Kallie | Black or African American | Cis-female/Cis-Woman | Health Promotion | Second |
| Adam | Black or African American | Cis-male/Cis-Man | Exercise and Sports Science | Third |
| Miley | Black or African American | Cis-female/Cis-Woman | Biomedical physiology and Psychology | First |
| Hanna | Black or African American | Cis-female/Cis-Woman | Biology | Fifth |
| Amelia | Black or African American | Cis-female/Cis-Woman | Biomedical physiology | First |
| Nicole | Black or African American | Cis-female/Cis-Woman | Public Health | First |
| Jayla | Black or African American | Cis-female/Cis-Woman | Pharmaceutical sciences | Fourth |
| James | Black or African American | Cis-male/Cis-Man | Agricultural Leadership, Education, and Communication | Sixth |
| Spencer | Black or African American | Cis-female/Cis-Woman | Chemistry | Second |
| Mary | Black or African American | Cis-female/Cis-Woman | Biomedical Physiology | First |

### Data Collection

The study was advertised via flyers across the institution, shared via science majors listservs, and emails/flyers shared by instructors who use active learning. Participants interested in the study were asked to fill out a survey through Qualtrics to ensure they met requirements. The survey provides a definition for peer discussion in active learning to help the participants to identify relevant courses to list (while we asked for course number, we did not ask for instructors). The survey also asked for participant demographics such as race/ethnicity, gender, sexuality, disability, first-generation status, as well as year in school, major, and high school.

#### Semi-structured interview protocol development

The interview protocol was piloted in Fall 2023. We interviewed 7 participants of color during our pilot study over via Zoom or in person and were compensated for their time with a $35 gift card. The protocol was modified several times for clarity and to better attend to our research questions. Questions were also added as ideas emerged. For example, it became clear in the first few pilot interviews that the demographic makeup of the high school the participants attended were related to how they made sense of their interpersonal interactions with White peers in their AL courses. Therefore, we added a question about how their interpersonal experiences with high school peers were similar or different to those at the institution. We also added more probing questions in case participants were not detailed in their responses.

Our interview protocol (see supplementary materials) included questions such as, “When you are interacting with peers in your science courses, are you ever reminded of any of your identities? “Please give me some examples of negative/positive interactions with classmates in science courses that used active learning”, “In the survey, you identified as [list of identities]. How, if at all, do feel that one or more of these identities played a role in your interactions?”. We proceeded to ask more specific questions about the experience, the demographic makeup of the group members or peers who contributed to the experience, and about how and with whom they make sense of or process the experience. We also asked about the structures put in place by instructors (faculty or TAs) to support peer discussions (e.g., group norms, group roles, explicit guidance or other salient practices for the participants). Finally, we asked “If you could give three pieces of advice for your course instructors to help improve peer discussions what would those be?”

#### Interviews

Semi-structured interviews were conducted in person or via Zoom depending on the participant’s availability and preference. The interviews were audio-recorded to allow the interviewer to focus on the participant’s responses. Participants read a definition of peer discussion in active learning prior to their interview to remind them of how this study is defining the term and centering peer discussions (see supplementary materials). All interviewers with Black participants were conducted by Black students (H.N., T.S., J.R.).

#### Field and Analytic Memos

Interviewers wrote post-interview memos (i.e., field notes) to document ideas and relevant details about the interview including non-verbal cues made by the participant, issues to consider during data analysis, and reflections on whether and how to improve the interview protocol or any portion of the methodology.

Analytic memos were also completed throughout the iterative coding process. These memos were used to document the researchers’ initial thoughts about data and later discuss common patterns that they were noticing amongst the participants. These patterns led to the development of themes. At least two researchers wrote analytical memos after the initial coding of each transcript. Subsequent analytical memos were written after the codebook was fully developed, but in summary format for a more comparative analysis, after re-coding every 6-8 transcripts.

### Data Analysis

A team of researchers (H.N., T.S., J.R., R.T., M.T., C.A., and T.R-T) conducted reflexive thematic analysis (RTA; Braun & Clarke, 2022). RTA is a process that involves developing, analyzing, and interpreting patterns across a dataset while critically reflecting on both your role as a researcher and the research process (Braun & Clarke, 2022). RTA is grounded in a subjectivist epistemology, which asserts that as our participants have knowledge within them and they are co-producers of the research findings, which is based on their understanding of their lived realities and are dialogically co-constructed with the researchers (Braun & Clarke, 2022). RTA involved six phases: data familiarization, coding, creating initial themes, refining and defining themes, as well as writing up our analytical processes and findings. It is important to note that these phases did not always occur in a sequential order, as each one requires several rounds of iteration and revising as themes are developed, codes and memos are revisited, and themes are eventually finalized.

While the codebook was being developed, the researchers noticed important trends that warranted further exploration. Namely, that the experiences of Black and African American participants shared specific commonalities that were not also consistently seen among Asian or Latiné participants. While Asian and Latiné students did report positive and negative racialized experiences, those tended to be more heterogenous. Therefore, considering the specificity of anti-Blackness for this sample, we decided to focus on Black students first and report on their findings separately with the care it deserves.

### Trustworthiness

We established trustworthiness for this project in a variety of ways. We conducted member checking interviews with 7 Black participants. Through this process, participants reviewed their interview transcripts and allowing them to add, remove, or clarify anything that they wish, we further shared a summary of our analysis and findings to confirm that we accurately captured their experiences (Braun & Clarke, 2022). We suspect that we were not able to recruit the entire sample of 21 students because we were recruiting for member checks in the summer. However, 1/3 of the sample allowed for additional data triangulation that we believed strengthened the trustworthiness of our findings. By coding to consensus, multiple researchers were able to check each other’s analysis and ultimately come to an agreement over what the data may mean (Braun & Clarke, 2022). Peer debriefing with researchers and colleagues who also engage in qualitative work and critical qualitative work was used to check for different perspectives, errors or bias, and finalize themes. To ensure transparency, which is a vital component of qualitative trustworthiness, we engaged in reflexivity by continuously and critically interrogating how our own subjectivity, values, and social positioning actively shaped the co-construction of our analytical insights (Braun & Clarke, 2022). Having previously detailed our epistemological and theoretical groundings, we outline our positionalities below, inviting readers to contextualize our interpretative lens and draw their own conclusions regarding the credibility of our findings (King et al., 2023).

### Researcher Positionality

Throughout this study–– from its conception to its implementation–– we remained highly attuned to how our individual and collective backgrounds shaped the research process. While we identify as racially minoritized people (Black and Latiné) with lived experiences of exclusion and inclusion in STEM, there is a robust diversity in our team across social identities (e.g., ethnicity, gender, neurodivergence, nationality, religion, and socioeconomic status) and professional roles (e.g., undergraduate researchers, graduate students, and a professor). Because we share similar science backgrounds and institutional contexts as our participants, our intersectional lived experiences deeply informed how we designed and executed this work. This rich array of standpoints enabled us to build affirming environments for the students we interviewed. At the same time, we worked to remain aware of the different levels of power and privilege we each wielded in our respective positions.

The prominent presence of Black researchers on our team afforded us a deep, culturally grounded understanding of the participants’ narratives. Many of us share distinct overlapping identities with our participants, such as navigating neurodivergence, being first-generation college students, or immigrating to the United States. While these shared positionalities allowed for authentic connection, we also recognized that each participant had their own unique intersectional experiences. To prevent our own personal histories from misrepresenting the participants’ truths, we leaned heavily on constant critical reflexivity, the foundational work of Black scholars, and consensus-building within our team to ensure analytical rigor and accurate representation.

Methodologically, we grounded our work in an asset-based, justice-centered framework that honors Black Students as the ultimate authorities on their own STEM experiences. By rejecting the hegemonic narrative that minoritized students must assimilate to succeed in STEM, our focus is on transforming educational environments so Black students can engage authentically and pursue their academic goals without having to contend with oppression.

## FINDINGS

Our findings illuminate how Black students experience negative and positive interactions during peer discussion in active learning science classrooms. These negative experiences were explored through the lenses of microaggressions while affirming experiences were documented using microaffirmations. While most participants connected these experiences to their social identities right away, others did not openly characterize their experiences as racialized until they were asked about how their social identities might have played a role in their peer-to-peer interactions or other related questions. Further, while participants also self-identified with a range of identities that included gender, race, ethnicity, LGBTQIA+ status, and generation status, for this sample, racial identity and gender appeared to be the most salient identities in peer interactions. Our findings also show that despite negative experiences, Black students still prefer active learning over solely lecturing and they had plenty of advice on how to improve it. We present these findings in the same order that the research questions were presented.

## RESEARCH QUESTION 1: NEGATIVE RACIALIZED EXPERIENCES

### Racial Microaggressions

We documented participants’ negative racialized experiences through various types of racial microaggressions. These racialized experiences were divided into 3 themes that represent the underlying exclusionary experiences of participants: 1) minimization and/or dismissal of intellectual contributions; 2) insensitive racialized remarks and behavior, and 3) racial and gendered discrimination. These themes help unpack the nuances of these experiences which, are all ultimately informed by macro-level racist and anti-Black narratives. We note that, while we assign themes and share specific quotes to exemplify them, microaggressions are layered and some of these experiences could fall into multiple themes. Quotes were edited for conciseness and clarity, removing filler words such as “like” or “hmm, etc. Table 2 describes the themes and provides extra instances of microaggressions experienced by students.

**Table 2.**
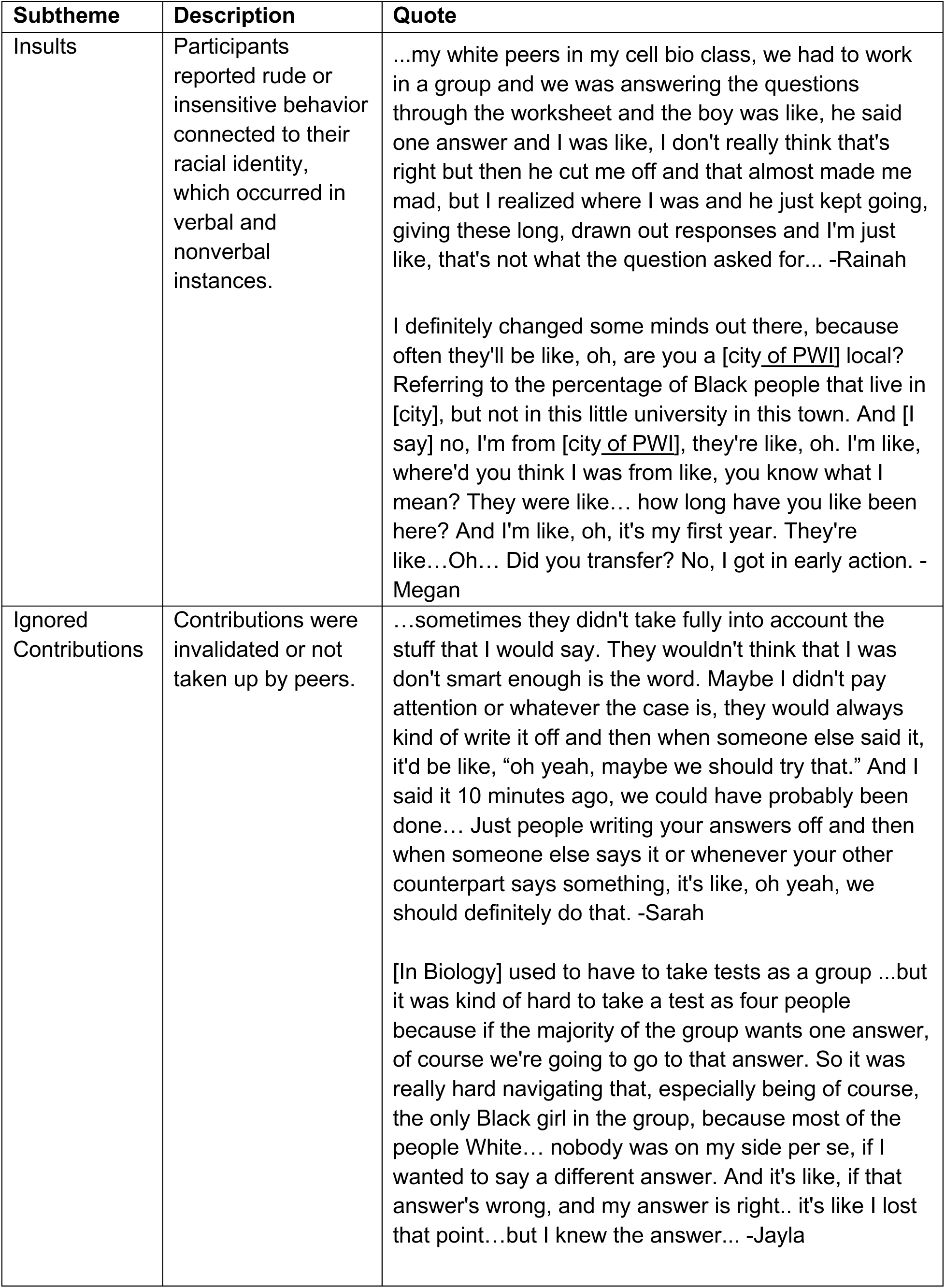
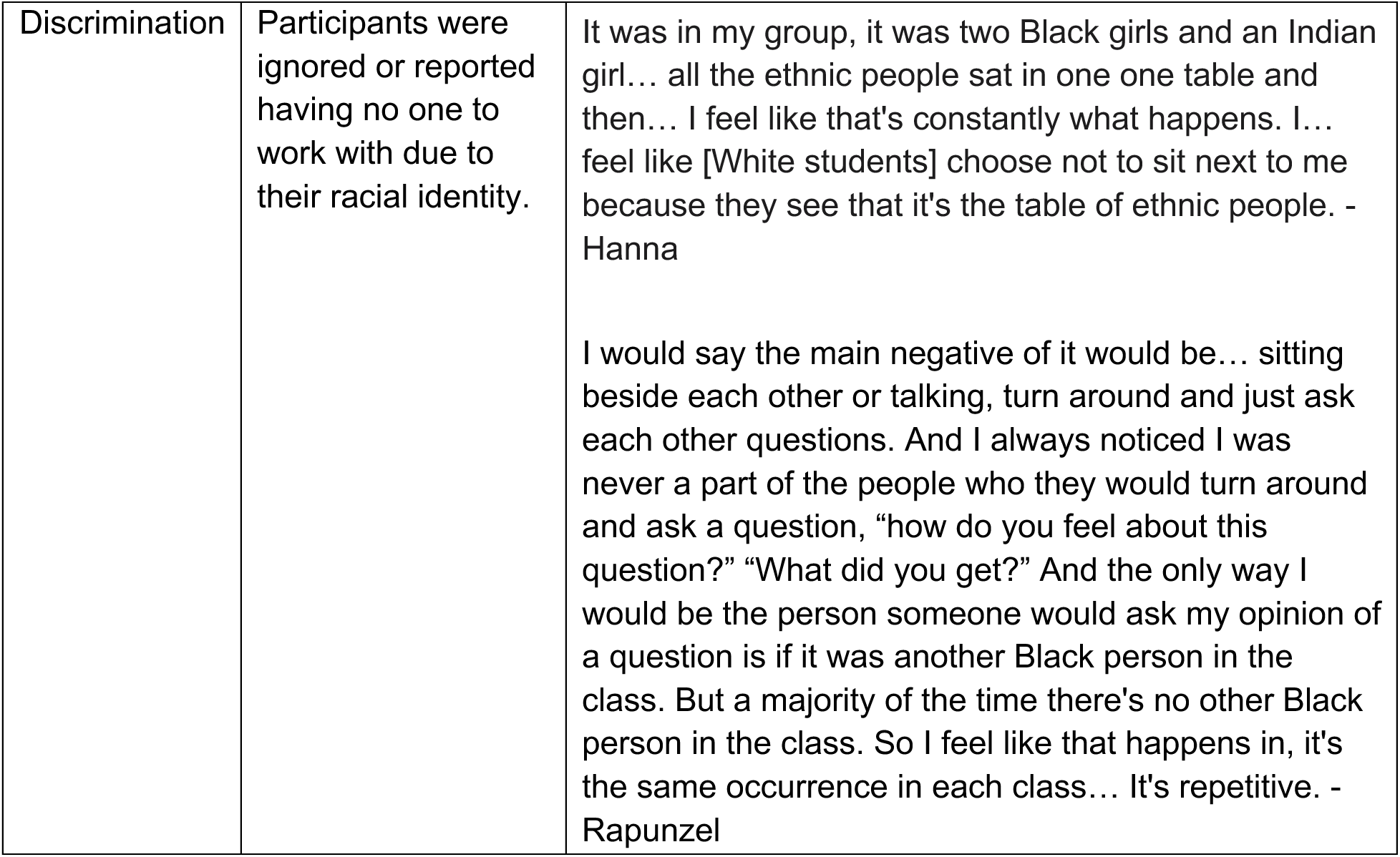
Exclusionary Racialized Interpersonal Experiences.

| **Subtheme** | **Description** | **Quote** |
| --- | --- | --- |
| Insults | Participants reported rude or insensitive behavior connected to their racial identity, which occurred in verbal and nonverbal instances. | ...my white peers in my cell bio class, we had to work in a group and we was answering the questions through the worksheet and the boy was like, he said one answer and I was like, I don't really think that's right but then he cut me off and that almost made me mad, but I realized where I was and he just kept going, giving these long, drawn out responses and I'm just like, that's not what the question asked for... -Rainah<br>I definitely changed some minds out there, because often they'll be like, oh, are you a [city of PWI] local? Referring to the percentage of Black people that live in [city], but not in this little university in this town. And [I say] no, I'm from [city of PWI], they're like, oh. I'm like, where'd you think I was from like, you know what I mean? They were like... how long have you like been here? And I'm like, oh, it's my first year. They're like...Oh... Did you transfer? No, I got in early action. -Megan |
| Ignored Contributions | Contributions were invalidated or not taken up by peers. | ...sometimes they didn't take fully into account the stuff that I would say. They wouldn't think that I was don't smart enough is the word. Maybe I didn't pay attention or whatever the case is, they would always kind of write it off and then when someone else said it, it'd be like, "oh yeah, maybe we should try that." And I said it 10 minutes ago, we could have probably been done... Just people writing your answers off and then when someone else says it or whenever your other counterpart says something, it's like, oh yeah, we should definitely do that. -Sarah<br>[In Biology] used to have to take tests as a group ...but it was kind of hard to take a test as four people because if the majority of the group wants one answer, of course we're going to go to that answer. So it was really hard navigating that, especially being of course, the only Black girl in the group, because most of the people White... nobody was on my side per se, if I wanted to say a different answer. And it's like, if that answer's wrong, and my answer is right.. it's like I lost that point...but I knew the answer... -Jayla |
| Discrimination | Participants were ignored or reported having no one to work with due to their racial identity. | It was in my group, it was two Black girls and an Indian girl... all the ethnic people sat in one one table and then... I feel like that's constantly what happens. I... feel like [White students] choose not to sit next to me because they see that it's the table of ethnic people. - Hanna<br>I would say the main negative of it would be... sitting beside each other or talking, turn around and just ask each other questions. And I always noticed I was never a part of the people who they would turn around and ask a question, "how do you feel about this question?" "What did you get?" And the only way I would be the person someone would ask my opinion of a question is if it was another Black person in the class. But a majority of the time there's no other Black person in the class. So I feel like that happens in, it's the same occurrence in each class... It's repetitive. - Rapunzel |

#### Theme 1.1: Minimization/dismissal of intellectual contributions

Participants’ negative experiences involved the minimization of their ideas and intellectual contributions. This can also be referred to as a microinvalidation, as these actions invalidate people of color’s experiences and knowledge. Participants shared how intellectual contributions such as offering answers to group assignments or providing feedback to peers were often scrutinized, dismissed, or ignored altogether. Black women specifically discussed that they believed these experiences occurred because of their status as both Black and Women–– a well-documented phenomenon described by Kimberly Crenshaw (1991). In STEM specifically, this “double bind” is an experience specific to women of color as they are forced to contend with others having a lowered expectations of them due to overlapping oppressions of racism and sexism, leading to the delegitimization of their credibility (Malcom et al., 1978). These microaggressions can discourage students from participating and further isolate the student from the rest of the group. Many participants discussed how their contributions were often met with skepticism. Megan shared:

> We were doing a group assignment for a worksheet, and I was getting answers here and there. They were getting answers here and there, but it seemed like every time I had an answer, and I would share with the group, it kind of be like, “how did you get that, or like, how did you do that?” And I take my time to explain it, but then I noticed that it was only reflected for me, but it wasn’t reflected for other members of the group.

Megan’s ideas and answers being constantly questioned reflects the work Black women must put in to be taken seriously. While having to explain one’s reasoning is part of active learning, Megan was the only one required to do this in her group. Thus, while she was trying to be helpful and build knowledge with peers, she was instead scrutinized and forced to prove the legitimacy of her answers. Rose talked about her experiences working with White men:

> In one of my courses, when I was in an all-girl group, one of them [said] “we won’t have to deal with white man syndrome”… sometimes it’s true because I’ve experienced that in other sessions. If I were to bring up something, guys in the class wouldn’t agree with me, and if later [I was found] to be [correct], it’s like that’s a different story as opposed to believing me. So maybe it does come from a variance of gender identities, but also racial as well…so many people can make different assumptions and then they’re definitely just not inclined to help you as much whatever the case may be.

Here, Rose unpacks how she often experienced interactions with overconfident and entitled white men who would think they knew the answer and assume that she did not because of their respective identities. Rose unpacked how it can make those who are racially minoritized more hesitant to speak up when confused, due to a fear of being judged harshly by those same White men and stereotyped them.

Black women explicitly talked about trying to collaborate with others but having their ideas dismissed or ignored. In response to, “Do you feel that any of your social identities affect your interactions with peers?”. Charity responded with:

> I feel like it’s related because I mean, naturally it’s just like some people might just think that oh, because I am a girl who’s African American, I might not be in a position where I can be as smart as them. So, say I’m in my Biology class, I’m talking to someone, collaborating with them like I’m sharing my opinions, but they’re just kind of like discarding it. Like “no, that’s wrong”, not even considering or even taking a moment to process it and give me a chance to see if it’s actually right. It’s just “no, that’s wrong” Yeah, it makes me feel like very dumb.

Due to her positionality as a Black woman, Charity’s contributions were not valued by her groupmates, and she knew this. Because her contributions were not taken up by her group, she began to internalize this in that moment doubted her own intellectual ability.

Invalidation and dismissal of ideas and contributions were the most common experienced microaggressions, as Black students shared various ways they were being dismissed by peers. As a result of their peers’ racialized and gendered assumptions, participants did not get an equitable chance to contribute to group work and were left questioning their own abilities.

#### Theme 1.2: Insensitive remarks and behavior

While not always related to discussions about class material, insensitive remarks and behavior were prominent in peer interactions in active learning classrooms. Also known as microinsults, these conveyed other types of deficit-oriented assumptions about participants such as being an affirmative action admit, participants’ supposed intentions or lack of work ethic. Microinsults were also conveyed via non-verbal behavior. ary explained her frustration with peers as they excluded her and another student. She did her best to get the entire group to focus on the project but was met with whispers and laughter from the two White girls in her group:

> I’m just like, okay, we don’t have to be friends here. Can we just do the work?… Then when me or my other partner would ask a question, they’ll be rolling their eyes and I’m just like, okay…Now I don’t even want to work on it.

Nevertheless, she tried to get them back on track, sharing:

> …obviously I’m Black and then the other girl that they weren’t speaking with, I think she’s South Asian… And in this situation, it was very apparent to me because it’s like the way that you can just tell when someone’s looking at you and they just automatically categorize you as less than… don’t even categorize me as human. They don’t think I’m going to pull my weight… I said, “okay, guys, let’s split up the work, let’s figure it out.” Now it’s crickets. Nobody’s talking. So I said, “okay, fine. I’ll do the introduction.” Then one of the [White] girls was like, “oh, you just want the easy part.” I don’t think that she’s a raging racist I don’t know. I don’t care. As long as she’s not putting that on in my face because that’s going to be the problem. But it’s like slight comments that they make or little quips or little laughs they do with their friends. It’s like, why can’t we all four laugh? I love to laugh. So it’s like small microaggressions.

Mary knew her peers were acting on stereotypes based on her racial identity, so she takes initiative to disprove those stereotypes. Yet, she is still met with disparaging behavior, despite giving them the benefit of the doubt as she did not want to assume racist intentions. Similarly, Rose, shared an experience of multiple microinsults that slighted her intentions and work ethic:

> I’ve been in so many different situations…people moving on and seeing that I was lagging behind, but they kept going; and another instance where we were all trying to figure out the problem, I clearly was trying to ask them for help and seeing where they were because I definitely think I was able to follow along, but they just kept making excuses like, “oh, we’re working it out, it’s okay.” Then, once they worked it out [they’d] say “hey, we finished, do you want to copy it?” that’s not what I asked for initially. It’s *collaborative learning*. I wanted to be able to do it with a set of people and that wasn’t what I was given.

In this short description, Rose shared how her peers did not support her and left her behind, and how she was not allowed to help solve a problem although she tried to be included (also a microinvalidation). Finally, her peers insult her work ethic and stereotype her as lazy, assuming she only wants to copy the answers versus being an active part of learning process, all while assuming they knew what was best. This inequitable dynamic positions Rose as deficient and unable to contribute and affects Rose’s ability to benefit fully from active learning.

While peer interactions can be beneficial for students learning the course content, these conversations can also become personal and help build community and trust. Thus, peer interactions can support students to connect and feel a sense of belonging; however, it can also lead to the opposite if students are confronted with microaggressions. Megan shares how a conversation about personal interests quickly turned into a series of racialized misconceptions:

> …They were talking about a Taylor swift concert, and I [shared], “Oh, my best friend, she went to an Eras tour concert.” My best friend is South Korean, but they don’t know that. So I’m guessing they assumed that my best friend was Black… they’re like, “Oh, my gosh… she went to the Eras tour…how did she find out about Taylor Swift?” I’m like, “what? Tay has been around for a minute. What do you mean?” I showed them a picture, and they’re like “Oh! Yeah.” So, they asked me where I’m from, And I said [town] and they were like “Oh, my gosh, [town]! like that’s such a nice area… what school did you go to?” and I was like [private High School] and they’re like, “Oh, my gosh! Like, how did you get into that school?”… why its like, every time, like I kind of like bring in a new topic, you kind of just like insert the stigmas into it when we’re literally in a class setting like, why is it so interrogative?… they had no idea.

Megan tried to join a casual conversation with her peers but was met with hostile assumptions and microinsults. Her peer’s surprise that Megan’s best friend (who was assumed to be Black) knew of Taylor Swift or that Megan was from a high-income neighborhood came from racist assumptions. Furthermore, their surprise to learn that Megan attended a selective private high school, could indicate that she was assumed to be diversity/affirmative action admission. Megan picked up on this and connected it to the racial stereotypes her peers had about her, which ultimately can affect belonging and trust in group work.

Whether it was assumptions that a Black student only wanted answers instead of collaboration, or misconceptions about the circumstances of their enrollment, or other deficit expectations about Black students’ backgrounds, participants expressed frustration with how they were being treated. Regardless of their peers’ awareness of these microinsults, they signaled to the participants that this was not a welcoming environment, which can affect learning and sense of belonging.

#### Theme 1.3: Covert Discrimination

While all microaggressions are based on racist assumptions, some appear to be more intentional and can be called microassaults. These microassaults are the most overt form of microaggressions as it is clearly done due to race and it is harder for one to argue whether this was intentional. In this case, it was a bit more covert, since it did not involve a verbal exchange––but blatant avoidance and evasion. As such, this subtheme differs from dismissal because participants were not even given a chance to contribute an idea to be dismissed and were instead were avoided all together. This microaggression often occurred when students were told to “turn to a peer” and discuss a concept, prompt, or question with someone near them. Upon that prompt, Black participants were visually acknowledged and then instantly avoided. This dismissal often occurred via non-verbal actions. For example, Amelia shared:

> Well, there are some people where like they would [appear] they’d be confused on something, and I had just finished getting the answer. They’ll look back to ask like a random person, and because I’m a Black girl, they wouldn’t really ask me. They’ll ask like someone else like an Asian person or another like White person that looks like them. And I’m just like, well I mean, I could have helped if you wanted.

Amelia recognized she was being ignored based on her racialized and gendered identity which was what when her peers initially saw when, they turned around during a think-pair-share. Tasha, shared how students near talked across her as if she were invisible. Her peers clearly saw her but chose to not include her and actively talk across her. Tasha internalized this, deciding to not even try to join the conversation as she knew they didn’t wish to interact with her. Although nonverbal, this was a dehumanizing microassault described by several participants. Adam shared a similar experience:

> I feel like it’s just slight things you’ll notice. [Professors] be like, “turn to your partner” and then they’ll turn to someone else, and then you’ll just be like, “I’m right here man.”

Adam explained the avoidance he faced and the small social cues he picked up on when he had to work with peers from dominant backgrounds. These cues essentially communicated to Adam he was not the person they wanted to collaborate with. Similarly, Jayla also discussed how she received signals that her peers were not interested in working with her:

> Yeah, I mean in classes where we don’t necessarily have group work, but [we] turn to our neighbors [and talk]. I’m always the one, no neighbor to turn to, or I’m just sitting there and it’s like, oh, well, they’re turning away. You know what I’m saying? They’re turning away [from me].

Jayla shares how whenever it is time for a think-pair-share, she is left with no one to turn and talk to. She later explained how when she takes classes with friends everything is fine, yet when in she is the only one in new spaces and trying to communicate with others, she is excluded.

Lastly, Hanna shared how she had to get used to microinsults:

> I think that I’ve gotten used to it now… Freshman year I would say that it affected me more because it was a cultural shock. I came from a predominantly Black high school… so it was just seeing people that didn’t look like me… and then not being chosen for group work would make me even more insecure.

Upon arriving to the institution Hanna felt uneasy not seeing people who looked like her because she attended a high school with a majority Black student population. Peer interactions in active learning exacerbated this feeling of isolation and lack of belonging for her as her peers avoided working with her. Overtime she became accustomed to these exclusionary experiences, but she should not have to do so.

## RESEARCH QUESTION 2: RACIALIZED AFFIRMING EXPERIENCES

### Microaffirmations

When asked about affirming experiences during peer interactions in active learning, participants discussed developing friendships and community, recognizing each other’s experiences, and building knowledge with one another. This also included having the opportunity to build friendships that outlasted the classroom, co-constructing knowledge with peers, and creating spaces where they could share advice and support each other’s success. We developed three themes that captured these racialized affirming experiences in peer discussion: 1) valued contributions; 2) feeling seen and supported; and 3) feeling safe and protected. However, a poignant pattern in these findings was that microaffirmations mostly occurred with other students of color–– and that they often had to be the ones who created the counterspaces to interact with each other to benefit from microaffirmations. Counterspaces are places (physical and otherwise) created for and (mostly) by people of color where one can authentically be themselves, feel affirmed in various experiences, unpack racialized experiences, avoid racial injustice, and create a sense of safety and inclusion (Ong et al., 2017). Table 3 describes the themes and provides extra examples of microaffirmations described by the participants.

**Table 3.**
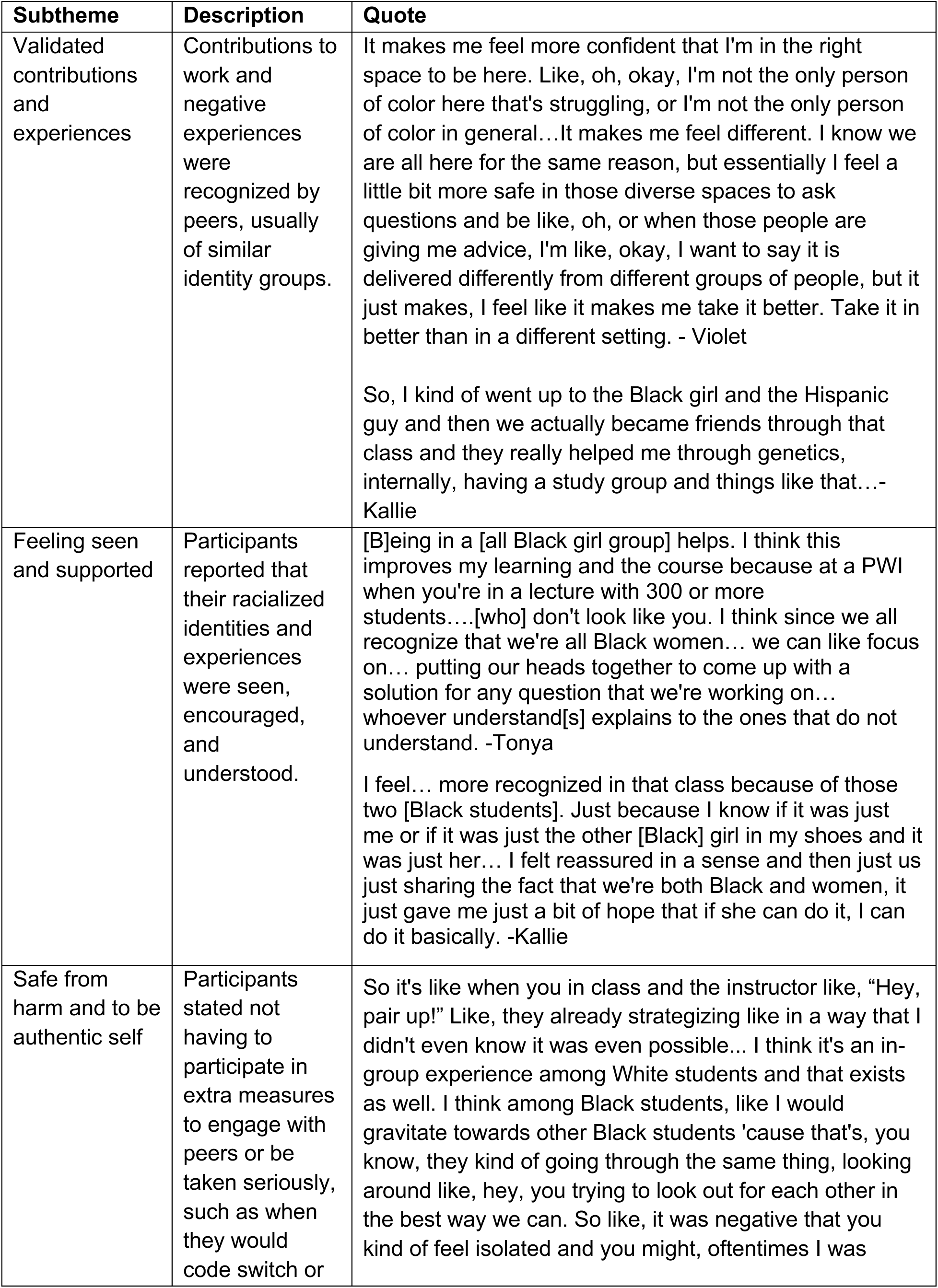
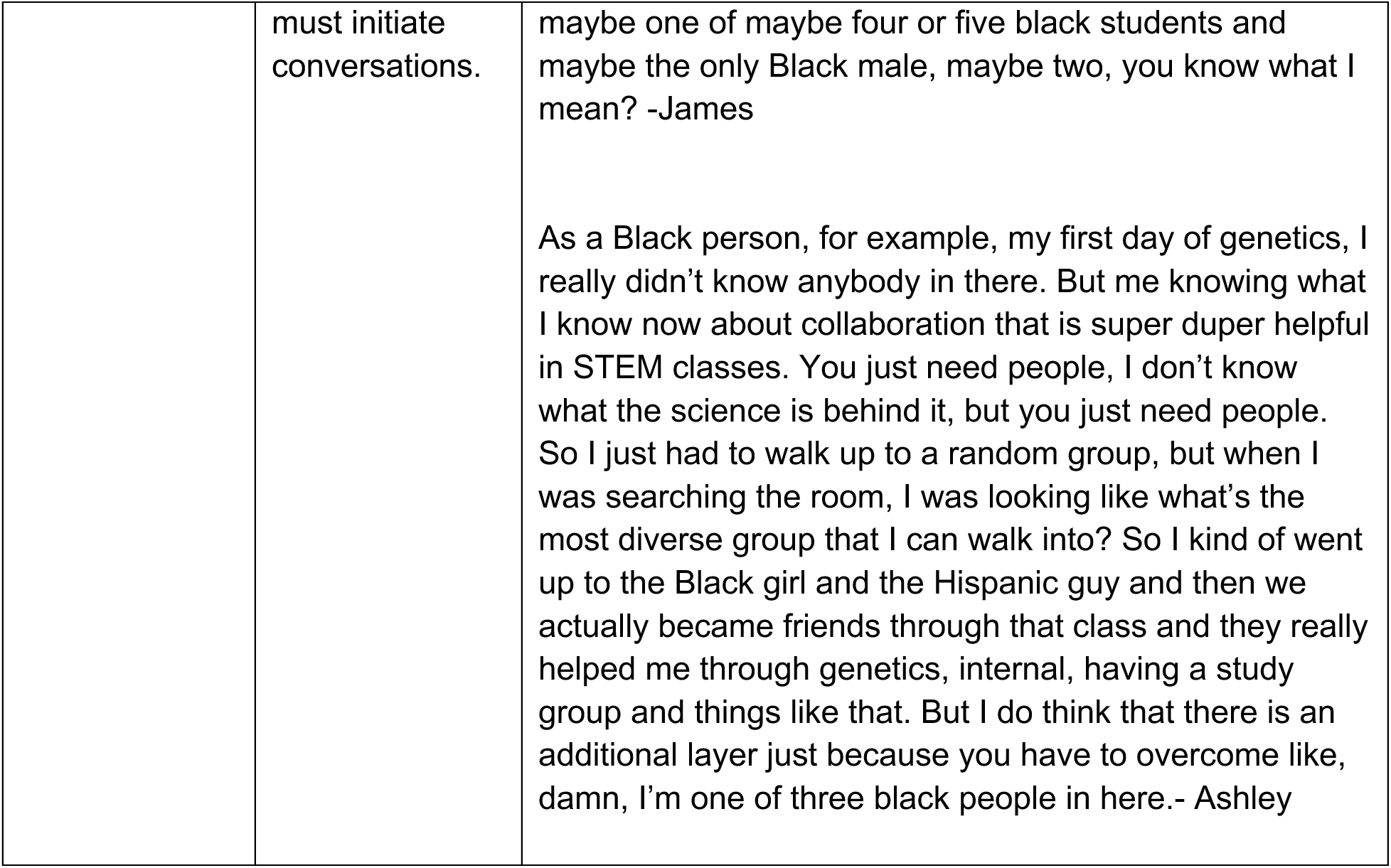
Affirming Racialized Interpersonal Experiences.

| **Subtheme** | **Description** | **Quote** |
| --- | --- | --- |
| Validated contributions and experiences | Contributions to work and negative experiences were recognized by peers, usually of similar identity groups. | It makes me feel more confident that I'm in the right space to be here. Like, oh, okay, I'm not the only person of color here that's struggling, or I'm not the only person of color in general...It makes me feel different. I know we are all here for the same reason, but essentially I feel a little bit more safe in those diverse spaces to ask questions and be like, oh, or when those people are giving me advice, I'm like, okay, I want to say it is delivered differently from different groups of people, but it just makes, I feel like it makes me take it better. Take it in better than in a different setting. - Violet<br>So, I kind of went up to the Black girl and the Hispanic guy and then we actually became friends through that class and they really helped me through genetics, internally, having a study group and things like that...- Kallie |
| Feeling seen and supported | Participants reported that their racialized identities and experiences were seen, encouraged, and understood. | [B]eing in a [all Black girl group] helps. I think this improves my learning and the course because at a PWI when you're in a lecture with 300 or more students....[who] don't look like you. I think since we all recognize that we're all Black women... we can like focus on... putting our heads together to come up with a solution for any question that we're working on... whoever understand[s] explains to the ones that do not understand. -Tonya<br>I feel... more recognized in that class because of those two [Black students]. Just because I know if it was just me or if it was just the other [Black] girl in my shoes and it was just her... I felt reassured in a sense and then just us just sharing the fact that we're both Black and women, it just gave me just a bit of hope that if she can do it, I can do it basically. -Kallie |
| Safe from harm and to be authentic self | Participants stated not having to participate in extra measures to engage with peers or be taken seriously, such as when they would code switch or | So it's like when you in class and the instructor like, "Hey, pair up!" Like, they already strategizing like in a way that I didn't even know it was even possible... I think it's an in-group experience among White students and that exists as well. I think among Black students, like I would gravitate towards other Black students 'cause that's, you know, they kind of going through the same thing, looking around like, hey, you trying to look out for each other in the best way we can. So like, it was negative that you kind of feel isolated and you might, oftentimes I was |
|  | must initiate conversations. | maybe one of maybe four or five black students and maybe the only Black male, maybe two, you know what I mean? -James<br>As a Black person, for example, my first day of genetics, I really didn't know anybody in there. But me knowing what I know now about collaboration that is super duper helpful in STEM classes. You just need people, I don't know what the science is behind it, but you just need people. So I just had to walk up to a random group, but when I was searching the room, I was looking like what's the most diverse group that I can walk into? So I kind of went up to the Black girl and the Hispanic guy and then we actually became friends through that class and they really helped me through genetics, internal, having a study group and things like that. But I do think that there is an additional layer just because you have to overcome like, damn, I'm one of three black people in here.- Ashley |

#### Theme 2.1: Validated contributions and experiences

When asked to share affirming experiences during peer discussions in active learning many participants discussed situations when they felt their intellectual contributions were taken up, valued, and appreciated by their peers. Even moments of confusion felt validating when they were acknowledged and taken up. Students reflected on these microvalidations by describing how they were able to build knowledge with one another, felt affirmed through verbal and nonverbal cues, and had respectful and productive conversations. These experiences also led to the development of study groups and friendships that became vital to the student’s experience in active learning across different courses.

Spencer talked how working with other peers of color led to recognition of her contributions. Small cues of validation, both verbal and nonverbal, helped her feel more comfortable in sharing ideas:

> …So a lot of the times…[I] said something, no one else heard it, and then someone else repeated what I said and all of a sudden people could hear them… but one of the classmates, we kind of got to know each other more was an Asian girl and she looked at me and acknowledged that she heard what I said…it was a little nice later on in the semester as I got to know [her] more, she would look at me and be like, “yeah, you were right when you shared the things”… “yeah, I heard you. I know you said what you said.” So that was nice.

Spencer appreciated how her classmate affirmed her contribution, even when they are the only ones to hear it. While these may seem like small interactions, they essentially validated Spencer’s contributions. This was even more meaningful for Spencer in particular, as she was a transfer student who was new to the institution at the time.

Being able to talk through confusion is one of the goals of peer discussion in active learning. However, that becomes difficult given the exclusionary experiences we previously described. Under those circumstances, being able to admit they are confused, articulate tentative explanations without judgement, and help each other understand the concepts felt extremely validating. Miley described such a validating experience when working with another Black student:

> In Chemistry, when our teacher put a discussion question up on the board or we are doing something and sometimes it’s just really confusing, I would explain it to my [Black] friend…if I get it, I’ll explain it to her. If she gets it, she explains it to me. So I’ll just be like, “wait, what does [the professor] mean?” And she’d be like, well, she’s trying to say this. Yesterday in chem, one of my friends helped me out. I was confused and she was explaining it to me, and that was really helpful.

In all, participants shared that working with peers who were also Black allowed for an implicit understanding of a shared experience navigating a PWI, which opened up space for participants to be vulnerable and be able to openly talk about their confusions in class in ways they did not feel possible with White students. Rainah shared:

> There’s not that many Black students here on campus…so when I do step into the classroom, I do find myself naturally gravitating towards people who do look like me and then it is just refreshing because not only are they a student here too, a minority here at [University] but we’re still navigating this big campus and still having to take classes here and we’re on the same page. They’re like “Oh yeah, I don’t know what the professor is talking about” … versus my White peers, they go, “oh yeah, I know [the content]. I’m like, somebody’s lying, but I’m going to let you have it. But yeah, that sense of familiarity and comfortability is reassuring.”

Students also pointed to the fact that science classrooms can be competitive environments that make it difficult for students to collaborate with others, and perhaps harder to admit when content is confusing. Rainah unpacked how she felt she could be open about her struggles around her Black peers. This was different from her experiences with her White peers, as she felt that they often put up a front of mastering the content. Knowing that one isn’t alone and being able to commiserate over points of confusion can be just as validating and affirming as having ideas taken up by a group. It was important for Black students’ sense of belonging to have their capabilities and knowledge valued and validated, along with their confusion. Microvalidations also appeared to help students became more confident in their answers and contributions to group work overtime.

#### Theme 2.2: Feeling seen and supported

Participants also reported microaffirmations which helped them feel seen, supported, and legitimized their racialized experiences. These types of microaffirmations are know as microrecognition. For example, participants were able to share their anti-Black experiences with other Black students in their group and receive recognition that those racialized experiences were real. This recognition made them feel seen and supported. Commiseration was a common experience for Black participants, where they shared negative experiences, showed empathy, and allowed one another to process those experiences. Kallie unpacked how simply being grouped with other Black students in a course empowered them to support each other:

> …It was me, the previous Black classmate that I was speaking to and then we had befriended another guy, he was a Black guy and [it was] just us… and by the end of the semester we were all sitting together. So honestly just being able to see each other and knowing that we’re all going through the same thing, I feel like that’s really affirming and then especially we’re not necessarily saying that we’re in the same friend group, but us as just a whole and we’re trying to move on together. We have similar career paths. We know what each other’s going through and we try to help each other, [whether] it’s through the questions that the teacher shows on the board and wants us to work together to figure out, or recitations, or even just understanding the lab even though we go on separate days. I just feel like that was really positive experience to have and I’m glad that the class brought me together with them.

Kallie shared how by the end of the semester, she and her peers had created a small safe space in the classroom where they could commiserate over similar racialized experiences as well as learning content. These spaces were a vital for our participants persistence as recognition of their lived realities is important in countering the cognitive load a person of color has to experience when dealing with microaggressions. Mary discussed this:

> Yeah. I would say in Chem, when I wasn’t talking to people who made me feel bad about myself, a lot of times I sat with a group of girls. We were all Black and all of us are trying to go into medicine. So having that community was really great because it’s like we are all after a shared goal. We’re all a similar identity. I mean, we were all Black women, so that was something that we were able to bond over and support and uplift each other in our studies, which I really liked… Even if I wasn’t doing the best in the class, I could always turn to somebody and we would figure it out together and it would be great…

Mary stated the importance of having a community of peers who have similar goals and lived experiences, which counteracted the microaggressions that affected her confidence. Being a STEM major is particularly challenging and can be very demanding for anyone, but navigating the double bind made it even harder to find community. When Mary found that community, she was able to lean on them for support and feel seen and supported.

Charity talked about the difficulty of navigating a majority White classroom when she didn’t know anyone. Over time, she learned to look for other Black people in the classroom to help her to find a supportive community that would recognize her racialized exclusionary experienced as valid and be able to support her in unpacking them. She went on to build lasting friendships with these peers through those shared experiences of being a Black student at a PWI.

> …If I actually do not know anyone in the class, my first instinct is to like go sit beside the Black person, the first Black person I see… Usually, they’re also like experiencing the… same class. So, we would just talk about [the racialized experiences]. Like can you imagine what just happened? And [it is affirming] because we both are sharing the same experience.

Being seen and supported by peers allowed for the acknowledgement of their lived realities that included but were not limited to being the only Black student and subtle forms of exclusion and racism. Participants were aware racial microaggressions were happening to them (even if at times they did not call it as such) and having these recognized by peers was an important way to affirm their experiences and take away some of the emotional labor of trying to make sense or rationalize away a racist experience.

#### Theme 2.3: Feeling protected from racist interactions

Participants explicitly expressed various instances of feeling shielded from harmful racist behavior by being in community with other students of color in their active learning classrooms. Through microprotections, students avoided microaggressions, were able to bring their full authentic selves to peer discussions and engaged in a series of racially affirming experiences. These microprotections and other microaffirmations described in the findings where only possible because students actively created these informal, small counterspaces. They were created in active learning classrooms when Black students worked with other peers of color whom they felt they could trust. By working together, they could empower each other and protect each other from racial microaggressions by other peers. These relationships would often extend beyond a single course. Ashley shared:

> Literally one of my best friends, I met her through organic chemistry, and we just found out we’re in the same class together and we’re like, okay, you’re going to be my study partner. Literally after that we’ve taken every single class with each other and I feel like that’s really helped just affirm, okay, I have another Black girl doing the same thing as me and we’re going through this together and it’s hard, but we can conquer it type of thing. Just having a community is very helpful in overcoming some of those things and then being like, oh my gosh, you experienced that too? Did you feel like them being weird to you? Or not wanting to help you, how they help other people and just being like, I’m not unique in this experience and so I don’t have to feel alone type of thing.

Meeting her best friend in organic chemistry provided Ashley with a comrade with whom she could be herself and take other science courses with, protecting them from microaggressions or at least allowing them to commiserate on the experience together. Importantly, these instances allowed participants to let their guard down. Working with those who provided microprotections, helped them avoid exclusionary experiences and get what they needed from the active learning experience. For example, Sarah shared:

> Well, my anatomy class, there was one girl, we were the same ethnicity… I didn’t feel like I had to code switch or whatever the case is, but I would talk to her sometimes and just ask her any questions about anything. I would talk to someone that wouldn’t deem me as like, oh, you’re just not paying attention, or, oh, you don’t know what you’re talking about. I would ask someone that would want to explain it to me.

Sarah was used to being stereotyped and having her ideas and questions dismissed by other peers. When she worked with another peer of color in her Anatomy course, she discussed how she didn’t have to spend extra energy code switching or risk being stereotyped. She was then able to put her energy into better understanding the content, as her peer created a safe environment for her to do so.

Working with other students of color created an environment that protected Black students from the threat of being invalidated, belittled, or ignored. Rapunzel shared:

> …Just having someone who understands what you’re going through and has had the… exact same experiences. Just having someone like that to be around, it just puts you in a more comfortable space in an environment where you just normally like to be to yourself and not speaking to anybody.

Rapunzel further unpacks how, by having peers in group discussion who are also Black, her experiences were understood, and she could let her guard down. Knowing that someone had shared experiences of exclusion put her in a space that led her to know her experiences were real and she could unload them because she was now in space where she was seen.

This form of microaffirmation proved to be important for creating environments where each and every student could be authentic and share the knowledge that they own. Being shielded from microaggressions allowed participants to focus more on their work and actually build knowledge with their peers. Microaffirmations kept participants going and building community despite the negative encounters. All positive experiences discussed above occurred in the context of the peer discussions in the active learning classroom, however, this only happened because Black students took the lead in creating small spaces with other students of color where their racialized lived experiences were recognized, and their contributions validated.

## RQ 3: NAVIGATING INTERACTIONS WITH COMMUNITY CULTURAL WEALTH

### Theme 3.1 Navigating Racial Microaggressions with Community Cultural Wealth

Participants relied on their community cultural wealth to persist through racial microaggressions and bias. Community cultural wealth draws from CRT and highlights several cultural capitals that people of color possess. Cultural capitals are the assets, skills and abilities a person of color develops from their family, community, and lived experiences navigating racist systems (Yosso, 2005). They include navigational, linguistic, aspirational, social, resistant, and familial capital. Familial capital are the abilities and tools students come to institutions with from their family and culture environments (Yosso, 2005). Navigational capital refers to how students of color successfully navigate oppressive institutions not made with them in mind, such as higher education. Social capital refers to the peers and contacts that help students access resources, while linguistic capital includes language and communication skills used to succeed (Yosso, 2005). Aspirational capital keeps students motivated to pursue their hopes and dreams in the face of barriers. Finally, resistance capital allows people of color to resist racism and its barriers. First, we focus on Sarah who shared how she used linguistic capital to be heard in peer discussions:

> In class, I code switch all the time. If I don’t, it’s just not going to turn out too well. Yeah, I have to do it most times, especially in a group setting. They’re not going to understand or they’re going to pretend to not understand or they’re going to write it off as some uneducated person. So, I’m like, okay, well, we’re just going to be on, an even playing field.

If Sarah didn’t code switch, she noticed her peers were unable to understand her or at least they acted as if they couldn’t. Code switching is a skill utilized by students of color students to more easily communicate with peers from majority backgrounds (CITE). Essentially, Sarah would code switch to make working together easier for her group and to ensure that her ideas were heard and taken up. However, this is not fair for Sarah who cannot be her authentic self and therefore had to spend energy translating her words and ideas in a way that comforted of her peers. On the other hand, Ashley navigated the same experience differently:

> Obviously, [racial microaggressions that dismissed her intellectual ability] originated from racism and slavery days, but if I’m being for real, I don’t really code switch around White people. I stay the same. I like to make jokes in class; I like to cut up, you know what I’m saying?

Ashley connected racial microaggressions directly to macrolevel issues such as slavery and racism. She displayed her resistance capital by refusing to code switch to make her White peers conformable, because she felt it diminished part of her personality.

When it came to finding someone to work with, participants often expressed that they had initiate the peer interactions themselves, as well as sustain it. This was despite picking up on cues that their partners did not want to collaborate. Hanna talked shared:

> I think that I’m pretty outgoing… I feel like I will talk to people, say I have a question or like he says to turn and talk to somebody. I feel like I’m always the one that’s making the first step, or I have to break the ice, or I have to talk to my partner, like they’re not just gonna talk to me… I also have gotten that I have like a RBF, which I don’t agree with. I feel like that is something with race as well…

Hanna used her navigational capital by taking on the extra effort to initiate think-pair-shares with her partners. She went on to discuss how she had been told that she has an unfriendly face (RBF). Hanna also displayed resistance capital as she is pushed back against a dominant narrative that Black women are unapproachable and mean.

Some participants discussed how their predominantly white high schools prepared them for navigating the predominantly white university and its classrooms. Charity shared how she developed the skills of proving herself to others back then:

> …So definitely like from high school, I already like got the mindset to not really like prove to myself to peers, but I guess show everyone that these assumptions you have aren’t true… like a Black woman can equally do really good in these more challenging classes and obviously I brought the same mindset over here [to the university]…I’m gonna finish college. [B]eing a first gen student, I’m gonna graduate, I’m gonna do good in a lot of my classes. That helped me to persevere in these situations… It encouraged me to also seek other Black students I have in my classes… we share the same goal of proving to society that this is not how it should be. It doesn’t need to be this way.

Charity had already developed skills to navigate a PWI. She knew that her peers would doubt her, so it was on her to reaffirm her identity and capabilities and find other Black students helped her do that. Charity utilizes navigational, social, and resistance capitals to locate community, reaffirm herself, and push back against racist narratives that inform interpersonal interactions in the classroom.

Participants shared how they used their skills and assets they developed from their lived experiences to navigate a racist environment. Some of these involved using methods of stereotype management, such as code switching and making oneself “approachable” by White standards. While all Black students, regardless of high school background, came to college already possessing community cultural wealth, the transition to a predominantly white university felt a bit smoother for those who had attended a majority white high school. Participants who had attended predominantly Black high schools, shared that the transition to that space was challenging and that they experienced a sharp culture shock. However, they quickly leaned in their navigational capital and were able to acclimate to the PWI.

All participants shared a myriad of ways they leveraged their various capitals to persist and resist in active learning classrooms and beyond. This also manifested in how they built affirming relationships which we unpack this in the next section.

### Theme 3.2: Leveraging Existing Counterspaces with Community Cultural Wealth

Community cultural wealth was also leveraged throughout participants’ positive experiences. In fact, their ability to create counterspaces in active learning classrooms speaks directly to the several cultural capitals they possess. As we discussed, most of our participants shared the importance of creating informal counterspaces in the classroom. Here further nuance how they navigated their courses by engaging with a counterspace outside their classroom. This large but informal counterspace was known as “Black [University]”. Black participants shared how leveraged Black University to make choices about the courses they chose to enroll in, to find other students who were in the same major and could take classes together, as well as community to commiserate with when they could not find other students of color in their own classrooms. Black University helped students to foster their resistance, aspirational, navigational, and social capitals. It did not only provide students with the opportunity to be in community but also create opportunities for sharing advice, resources, and mentors. Jayla reflected on Black University helped her to develop her in class counterspaces:

> …I had a lot of classes with my besties, and I feel like we really were each other’s support system… we studied together. We sat together in all the classes we had…I feel like that’s how we pretty much got through all our classes together. If one didn’t know something, we tried to help each other out. We all went to office hours together to try to understand stuff together and it worked for us… Definitely being Black and being exposed to Black University as soon as I got to campus allowed me to meet the Black people that were in my field and going to be taking classes with me.

Kallie also talks about how Black University proved to be an important tool for navigating the institution and their classrooms. She talks about how Black University connected her with some of her fellow classmates via social media:

> Being a Black woman and finding another Black woman or Black man in the same field, especially in Chemistry [is challenging]… Black university had their [Instagram account]… so I had already known this girl beforehand [as] we were mutuals on Instagram… I went up to her the first time we had chemistry class together… I knew she was friends with some of my friends, but we really never spoke. I reintroduced myself and then we just started sitting together. So just us having that little source for us…just us being on the Instagram page together [helped].

For students who were unable to find other students of color in their majors or who could take the same courses together, the peers that participants met through Black University provided them with the space to vent about such experiences, as Rapunzel shared:

> I mean, you just had to brush it [being avoided] off, but when it did bother me, I did have [Black] friends that I would go to and be like, yeah, this happened. A student was talking to me and stuff, I didn’t know what was going on and then I like tell a friend about this and they’d be like, “yeah, same thing happened to me, but honestly it’s okay. You know your worth, you’re good.”

Finally, Mary discussed using her navigational and resistance capital to counter feelings of isolation in her courses and peer discussions. She and other Black peers refused to let their racial minority status affect how they saw themselves in the classroom and instead used it as a source of empowerment:

> With being Black, I would say it would create comradery because we all recognize the struggles that we have faced and we’ll probably continue to face. So instead of using that as let’s all sit around and have a pity party for ourselves… let’s use it to uplift ourselves and be like, okay, let’s be Black excellence. Let’s be women in STEM… So, we’re able to be like, okay, you know what? We’re in a field that doesn’t have many people that look like us. We all want to make it, so let’s all help each other get there.

Here Mary displayed several capitals, from resistance and social, to navigational, and aspirational. She acknowledged how her racialized and gendered identities in STEM create barriers for her and her peers, but they also opportunities for them to bond and empower themselves.

Students of color come to college with many skills and tools that they have developed with the support of their families, community, and culture. Black students leveraged those skills and assets to navigate the institution and their classrooms. This was no different for how they navigated their positive and negative experiences in science active learning classrooms. These strengths were vital in helping students to persist through and resist micro-level systemic barriers and engender empowerment and community.

## RQ 4: PERSPECTIVES ABOUT ACTIVE LEARNING VS. LECTURE

Considering their overall experiences, participants were also asked if they believed that active learning was beneficial, and whether they preferred active learning over lecture. Except for one person, all participants in the sample reported that they preferred active learning and would NOT like to go back to a solely lecture environment. This is despite negative experiences and the extra labor they had to engage in to create their own opportunities for positive experiences in peer discussion. Nicole explained:

> Of course I feel like it’s good ’cause like sometimes I feel if it’s just too much listening from the teacher and there’s not like just a pause to just discuss what you’ve learned. Even if the group is not [the best] I feel like it’s just good to like step away from the lecturing for a couple seconds and just discuss it, soak [it in] and at least help us like think to ourselves regardless of how the group is like, just think to ourselves about what we just learned. So, I think it’s definitely been beneficial.- Nicole

This is strong evidence that students indeed believed that active learning was beneficial. In Sarah’s own words “It is beneficial, when done correctly.” Below, we briefly discuss two overarching themes that represent most of the participants’ responses to these questions. Table 4 summarizes the themes and provides more examples of student’s perspective about active learning.

**Table 4.** Perceived benefits and missed opportunities of active learning.

| **Subtheme** | **Description** | **Quote** |
| --- | --- | --- |
| Perceived benefits | Participants described active learning as enhancing learning, engagement, and peer connection. | Absolutely [I prefer active learning] It's an interactive environment. Even when you go into professional fields ... You're going to be talking to people with different backgrounds and interacting and bouncing ideas off each other. So it's good to have that, it reinforces the ideas more... because you're like, we talked about this and this and this. You know, it kind of helps cement that. Even in the real world, like, you're not gonna be doing things by yourself. -Spencer<br>...as teachers you're teaching the same materials over and over again, so you just know it and just spit out the information for us to understand. But then when we're put in smaller groups, it's like first, you're working with a smaller group of people... you're no longer as shy or feel like you're gonna be like judged by... the bigger class as a whole. And also...getting explanations from students, it just like feels better ...They explain it better knowing from student to student... like you're on the same level. So, they understand the thought process and it's just the little stuff like giving you easier ways to learn the material... -Charity |
| Missed opportunities | Participants emphasized that active learning effectiveness depended on implementation. | Maybe if we like, were able to like to get more time and like the teacher can explain it briefly after we've done the question so we can understand what just happened instead of going through so fast. Because I feel like most of like the active learning in classrooms, like whether it's with groups or with like square cap or like any clicker questions, it just goes by so fast. Like you can't keep up with what you don't know and what you know. Or if you know you don't know it, you can't go back to it 'cause you're already like five questions in at that point. -Nicole<br>They really don't guide us at all... Now definitely a little bit more guidance as to how things should flow in some of our group work. There's a lot of group projects that they definitely outline the tasks and things that need to be done... but before I feel like we were kind of thrown to the wolves. You just work together. -Jayla |

### Theme 4.1: Benefits of Active Learning

Students described active learning as beneficial for both learning and social engagement, highlighting its role in supporting academic success, creating more comfortable learning environments, and fostering a sense of community among peers, and preparing them for the workforce. Rose characterized active learning as a critical component of academic success by emphasizing the importance of collaboration, peer interaction, and problem solving in the learning process:

> I don’t think I’ve ever gone a year in all of my years of schooling and have been successful without [peer discussion]. I think that there’s always going to be a time where you need to either ask for help or really work with others in order to understand something or to work towards a common goal. Some situations you can work independently and still do amazing things, but I think sometimes with achievement it can be dependent or a lot more helpful with someone else or with the team of people that understand what needs to get done and may bring different things to the table, different ideas, different kind of innovative works or kind of understandings that you may not have thought of before but may help you within the next group of people that you transfer to.

Rose shared that active learning environments can foster deeper understanding by encouraging students to exchange and build on their diverse sets of knowledge, insights, and work collaboratively towards shared academic goals. Hanna emphasized another positive facet of active learning: creating a more comfortable and less judgmental learning environment compared to lecturing and getting to meet people who are different than her.

> I would still say overall I do think that it’s good to talk about stuff with a student rather than, just your professor. It is not the same ‘cause you might feel like you’re asking a dumb question or something, or they might move too fast. If you have another student, you could sit down with and go as slow as you want… [E]ven though most of the time I work with people of color, it also shows me [that I can] make more friends or… see who else is on campus.

Hanna described the ability to engage with other peers as reducing barriers to participation, in particular, the fear of asking “dumb questions” or struggling to keep pace in lecture settings. Hanna also noted that AL facilitated social connection, through interactions with other students of color as well as introducing her to students who are different. Similarly, other students described peer discussion and active learning as beneficial not only for learning, but also for fostering social connections and reducing feelings of isolation in the classroom. For these students, AL also contributed to relationship-building and a stronger sense of belonging within the university setting, as Sam described:

> I would say [it is beneficial] because I have formed positive connections through collaborative learning, and every time I’ve collabed with somebody, it has made me, juxtapose how I felt first walking into the room–– it’s all tight, nobody knows each other. But through collaborative learning, I feel like that tightness, that stiffness, that reservedness is just slowly brought down. And honestly, it fosters a sense of unity for me in those spaces. Walk into a room and it’s like, okay, everybody is definitely going through what I’m going through. They’re just trying to understand the material, trying to do well on this next test and then move forward. And I think collaborative learning has helped it not feel so individualized, but hey, we’re all moving through this together.

Sam positions collaborative learning as a key component for fostering a sense of belonging in the classroom. He describes an initial sense of discomfort and social distance upon entering the learning space, which is gradually eased through repeated collaborative interaction. Over time, active learning environments are framed as shifting classroom atmosphere from isolation to one of shared academic struggles and mutual understanding. Sam emphasizes a sense of unity, where peers are viewed collectively navigating similar academic experiences rather than competing.

### Theme 4.2: “Yes, but…” Missed Opportunities

Although many students acknowledged the benefits of active learning, they also emphasized that these benefits were highly dependent on how activities were implemented in the classroom, which makes sense, given the experiences described in the prior sections. Participants described ineffective active learning experiences as unstructured, forced, or lacking clear instructions and/or structure. Some also discussed when peer discussion or group work failed to promote meaningful group engagement or deeper critical thinking, with problem sets or activities that did not foster thoughtful discussion and just “busy work” that could be done independently. This suggests that students believe that active learning is most beneficial when intentionally facilitated, intellectually challenging, and clearly connected to learning objectives. For example, James shared:

> I think it has the capacity to [be beneficial], I think the way that it’s being done kind of haphazardly and maybe without intention is not appropriate.

Similarly, Miley describes how group discussion can be unproductive if not properly structured:

> In class, my teacher always gets us in a group talk. She puts that question out and [tells the] groups to talk about it. And no one ever wants to do it…everyone’s just sitting down there, staring at the board… Nothing ever productive comes out of that because it seems forced. Most times it’s just everyone saying, “oh yeah, this is what I think.” But no one elaborates on why they think that… We don’t really go in depth into why we think that because we are just doing what we are told to do… it’s not really that beneficial [when done in that way].

Miley shared how implementation matters and if the discussion feels forced and is not properly facilitated, the learning, if any, remains surface level and, as a result, the activity feels like a means of compliance rather than meaningful learning.

Overall, students described active learning as beneficial for learning and peer connection but emphasized that its effectiveness depends on how it is structured and facilitated in the classroom. Yet, they still prefer active learning over lecture.

## RQ5: FEEDBACK TO INSTRUCTORS

Given that out participants have a wealth of knowledge, skills, and lived experiences on the topic, we also asked them to share feedback on how instructors could improve active learning environments to make them more supportive and affirming. Participants’ answers centered around 3 themes: 1) class structure, 2) class climate, and 3) the need for explicit training in collaborative learning. It is also important to note that much of the students’ feedback stemmed from positive experiences they had in courses where instructors appeared to have implemented active learning in an equitable way and when race and other identities did not become as salient for our participants during peer discussions. Table 5 provides descriptions of the themes and additional example quotes from participants.

**Table 5.**
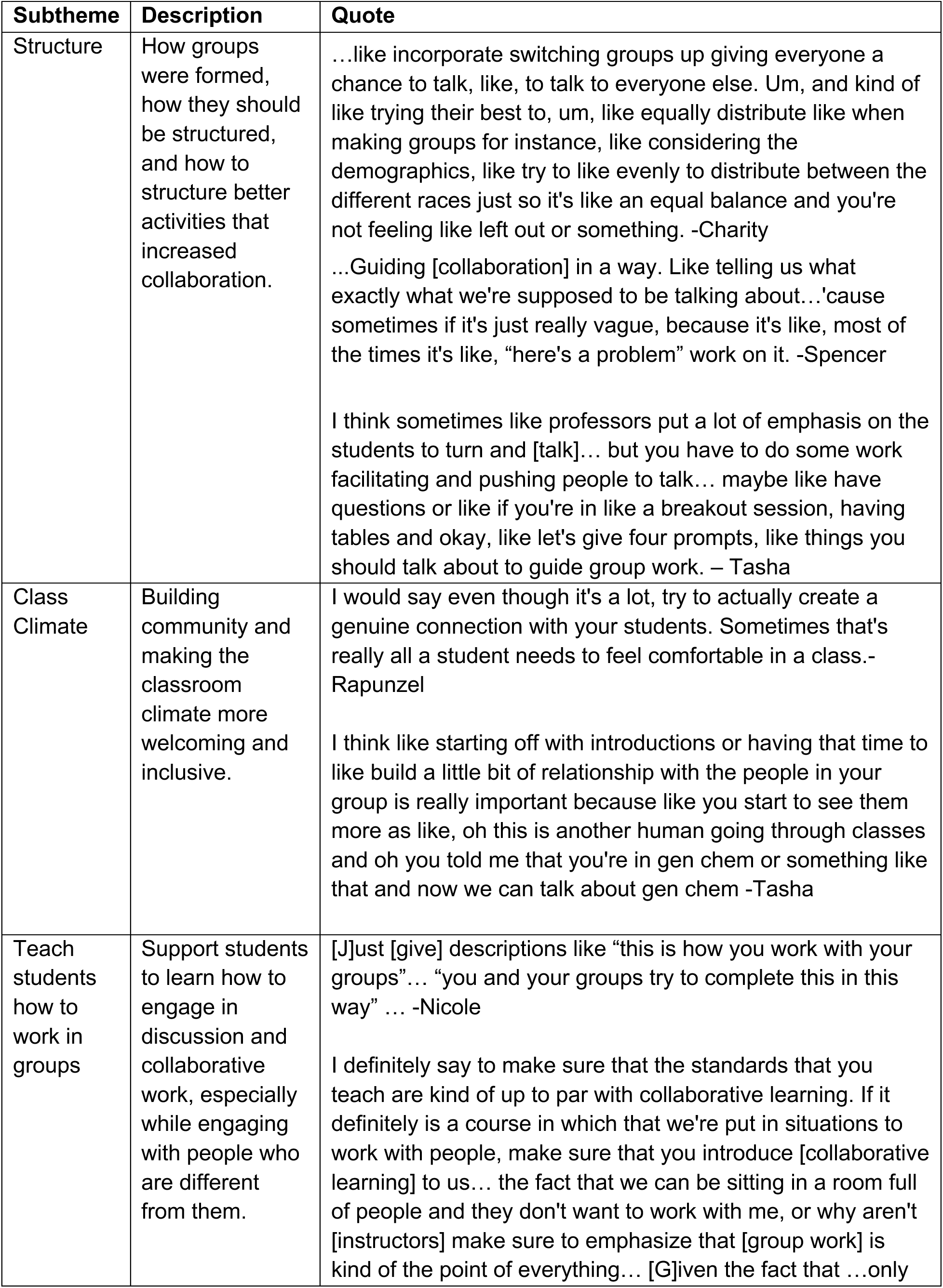
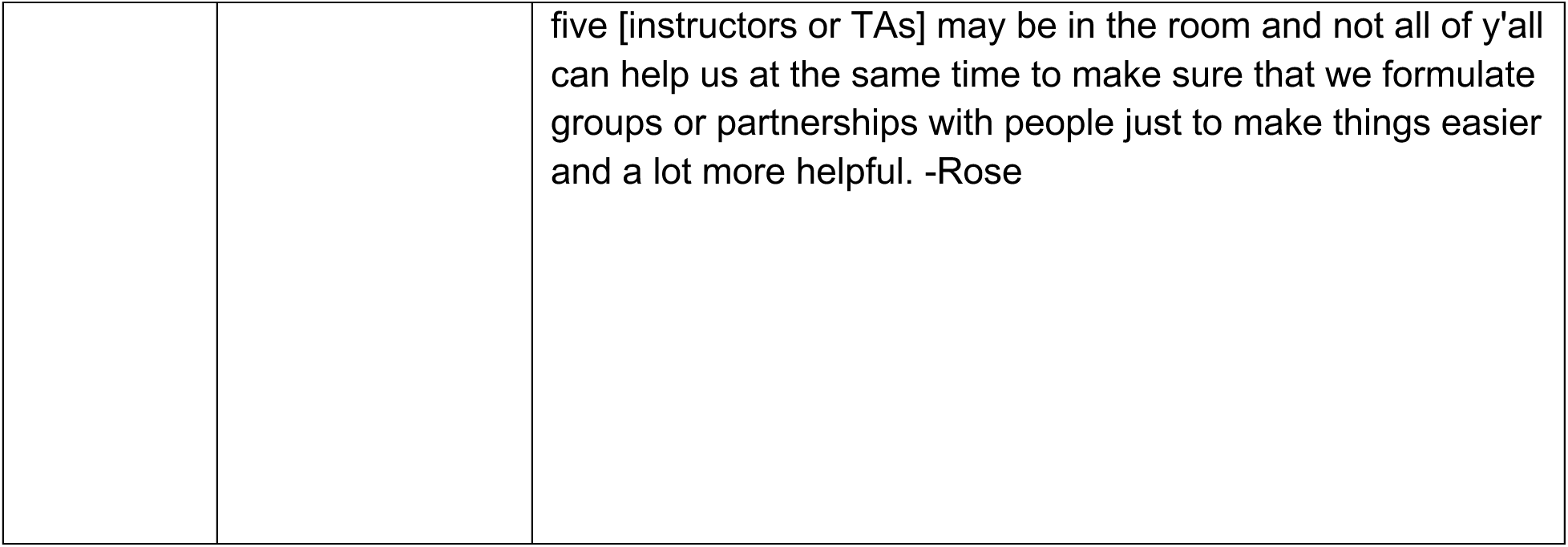
Feedback to instructors on how to improve interpersonal interactions.

| Subtheme | Description | Quote |
| --- | --- | --- |
| Structure | How groups were formed, how they should be structured, and how to structure better activities that increased collaboration. | ...like incorporate switching groups up giving everyone a chance to talk, like, to talk to everyone else. Um, and kind of like trying their best to, um, like equally distribute like when making groups for instance, like considering the demographics, like try to like evenly to distribute between the different races just so it's like an equal balance and you're not feeling like left out or something. -Charity<br>...Guiding [collaboration] in a way. Like telling us what exactly what we're supposed to be talking about...'cause sometimes if it's just really vague, because it's like, most of the times it's like, "here's a problem" work on it. -Spencer<br>I think sometimes like professors put a lot of emphasis on the students to turn and [talk]... but you have to do some work facilitating and pushing people to talk... maybe like have questions or like if you're in like a breakout session, having tables and okay, like let's give four prompts, like things you should talk about to guide group work. – Tasha |
| Class Climate | Building community and making the classroom climate more welcoming and inclusive. | I would say even though it's a lot, try to actually create a genuine connection with your students. Sometimes that's really all a student needs to feel comfortable in a class.- Rapunzel<br>I think like starting off with introductions or having that time to like build a little bit of relationship with the people in your group is really important because like you start to see them more as like, oh this is another human going through classes and oh you told me that you're in gen chem or something like that and now we can talk about gen chem -Tasha |
| Teach students how to work in groups | Support students to learn how to engage in discussion and collaborative work, especially while engaging with people who are different from them. | [J]ust [give] descriptions like "this is how you work with your groups"... "you and your groups try to complete this in this way" ... -Nicole<br>I definitely say to make sure that the standards that you teach are kind of up to par with collaborative learning. If it definitely is a course in which that we're put in situations to work with people, make sure that you introduce [collaborative learning] to us... the fact that we can be sitting in a room full of people and they don't want to work with me, or why aren't [instructors] make sure to emphasize that [group work] is kind of the point of everything... [G]iven the fact that ...only |
|  |  | five [instructors or TAs] may be in the room and not all of y'all can help us at the same time to make sure that we formulate groups or partnerships with people just to make things easier and a lot more helpful. -Rose |

### Theme 5.1: Class Structure

For participants, structure included both how groups were formed, how group roles were decided, whether groups should be permanent for the semester–– and also how the actual learning activity was structured. Structure had much to do with what students perceived was best for their learning, while also considering what would mitigate racial microaggressions. Many cherished the idea and benefits of being part of a diverse group. When it came to how to decide how to form groups for peer discussion and group work, some students preferred to be randomly assigned, while the others wanted to pick their own groups. Both categories of participants preferred having the opportunity to shuffle groups at some point in the class or at least have an opportunity to transition out of groups if conditions warranted. Participants who preferred being allowed to select their own group tended to describe experiences where there were more students of color in the classroom, or when at least they felt comfortable going up to the few students of color when it was time for form groups. Students who preferred random group assignment, tended to have been enrolled in classes where they were often “the only one.” Randomization would therefore take away the tension of having to find their own groups and risk microaggressions, while providing them with opportunities to get to know different people and learn from them. James explained his view:

> I think professors should be more intentional about how people are grouped and maybe preventing the isolation… or like [provide] a continuity of like, I’m always going to work with these people on any [think-pair-share].

Students from both categories also discussed how picking their own groups and not switching would make the collaboration experience smoother. Mary shared:

> Maybe if you allow the students to pick their groups, I know that that can be really difficult because there’s a billion people in the class and you don’t really know how to split it up, and it makes it a lot more work for the professor. But I think it would… be a lot less painful because if I was working with the people that I sat around, we’ve all known each other all semester. I have a good idea of how they work, how they study. I continue to sit by them because I actually like them. So why not? Let’s just wrap it up and do a group work together.

Mary already enjoyed working with her peers and would rather continue to build that comradery with them rather than switch groups. The common trend was that their classes did have at least some people of color and if they are just allowed to sit together, it would not be a big issue.

A few participants, like Hanna, proposed the flip-side of this strategy:

> I don’t know if I would like this honestly, but it might help somebody. I would kind of do randomized or like “sit with someone new” this day or something like that because I feel like that would have forced me to maybe work with a man… Like work with a white woman or just someone who’s different from me. So, I think forcing randomized or just switching up sometimes at least, like you can have your favorite, but some days let’s do random seating.

Finally, beyond group composition, many participants talked about how the structure of group activities and assignments themselves needed to be improved. Jayla highlighted how the nature of the work can affect whether group members choose to work together:

> If they’re going to do group interactions, definitely make it a lot more structured. Give us what you want us to do within those groups and how you want us to do it because a lot of those collaborative assignments aren’t collaboration for real. Either one person does it or we all do it on one doc. We don’t really talk to each other about it and if you’re going to do group work, make it interactive work. If it’s something that I can just do on my computer and turn in, we’re not going to talk for real. Pretty much the same as the second one make it so we need to meet up and actually do the work together.

Jayla details how many of the in-class group work she has given only required that the worksheet be completed, not necessarily encouraging the groups to co-construct knowledge, or collectively grapple with the material.

Thus, more equitable structure for students involved improving how to pick groups, the diversity of the group, and the facilitation of discussion and assignments. Students had varying ideas on how groups should be formed but all agreed that diversity in groups was important so that everyone’s ideas were heard and no one was left out. Along with diversity and group formation, they discussed intentional group work design, so that assignments required students to trully collaborate.

### Theme 5.2: Class Climate

Our participants also discussed the importance of establishing a welcoming class climate by building community with their peers and instructors, engaging in practices that discourages competition, and humanizing the classroom. When students felt more comfortable with their groups, it was easier to talk about content and break away from shyness or fears of being negatively judged. They believed that classroom climate began with the instructor. Rapunzel shared:

> I would say even though it’s a lot, try to actually create a genuine connection with your students. Sometimes that’s really all a student needs to feel comfortable in a class.

Adam added to this train of thought by sharing:

I really do enjoy learning more about my professors and why they chose their field of study. They might be talking about the subject and I’m just like, I don’t care. I don’t care about Social Science. This is not the thing I’m interested in but if they explain, yeah, I chose this because X, Y, Z, and hearing their background is kind of interesting to me, it’d be like, okay, I can kind of get behind this or also just, it builds the relationship because you get to know more about your student and you make the student feel like they’re valued because sometimes you just don’t feel valued. [T]eacher-student connection is important.

Taking time to share a personal piece of their life and humanize the classroom helped students see professors as people and not just authority figures, creating a warmer climate that can begin to foster a community, which was important to students given they were expected to engage in peer discussions and group work on a daily basis.

Yet despite how important having community felt to students, they surfaced an important contradiction between progressive active learning teaching practices co-existing with traditional grading practices that foster competition (i.e., curving). Ashley shared a story that highlighted her experience with how this common exclusionary assessment practice was at tension with creating a climate for true collaborative learning:

> …I feel like I’m not super competitive. I’m the type of person, if we all need to eat, let’s all eat. You know what I’m saying? So, when I got into STEM, I feel like I was very naïve of that, but then I realized people literally intentionally do things to make sure that you don’t get things. For example, I know you probably experienced this, but literally before test day for example, before you go into chem exam… People literally start being like, do you remember this? Do you remember this? And sometimes it don’t even be because they’re trying to learn it, it’s because they be trying to psych everybody else out… I know you hear that a lot like, oh, this was so easy I didn’t study…But I feel like in STEM, people do things like that. Like okay, they know it’s like a race from the get-go because of med school, any type of professional school is very competitive… and yeah, it’s just very competitive. It’s very hard finding people who will actually be willing to help you instead of competing with you.

Breaking down the competitive climate was something students often pointed to but didn’t know how to directly state in their advice to instructors because many did not recognize that curving grades was a choice professors made, as opposed to just as a “normal” part of STEM classrooms.

### Theme 5.3. Teach students how to work in groups

Similarly, Black students suggested that faculty learn how to facilitate group discussion, and peers learn how to communicate across cultural differences–– and that all students, including Black students–– needed support on how to work in groups. Since students are navigating new people, cultures, and environments (some for the very first time in college) they find it is important that everyone is guided in how to do this properly so that there are shared language and practices understood by everyone. Some participants alluded to the fact that instructors often assume students know how to work in groups. Sarah stated:

> I feel like teachers and students should know that not everyone knows how to work in a group. And I know that we’re all college students, early twenties. But I guess giving a brief rundown of what needs to happen in the group and making sure that those things are being [carried out]. Making sure that everyone’s ideas are heard, making sure, I guess just giving a rundown of what working in a group is like, because sometimes people, maybe they have worked in a group or didn’t know how to do it effectively, whatever the case is. So, I feel like that would probably be things that I would change.

Tasha discussed how she felt one of the simple things one of her professors emphasized helped to shape more inclusive interpersonal interactions:

> Like in my ecology class, she really made a point to be like, “turn to your neighbor and don’t be a jerk”. And I feel like when you emphasize not excluding people like that…someone is right there [don’t] act like you don’t notice them. Like include them.

This simple and direct reminder proved to be effective in Tasha’s experience. She felt it gave visibility to the students who are often overlooked. Tasha also discussed the need to tackle racialized exclusion more directly. However, like many other participants, despite sharing clearly racist experiences, she gave her White peers the benefit of the doubt appearing by saying that perhaps White people were not exposed to people of color in their hometowns and schools, which allowed for stereotypes to take hold. Tasha stated:

> Maybe it’s like they just haven’t been exposed to a lot of Black people. Like, because I know, like for example, like I’m from [majority Black town] and like I was the only person from my graduating class to come to [University] from my high school… Like if they haven’t had [exposure], and the amount of Black people up there is so slim. Like they really haven’t had those [cross cultural] experiences. So maybe they don’t [know].

As we have seen, many participants discussed experiencing microaggressions from White students. However, the vast majority did not assume their White peers had bad intentions. Like Tasha, many participants blamed the macro-level narratives and sociohistorical context for those problematic interactions and wished that White students would learn to be better. These experiences cleary speak to the need for active learning classrooms and institutions to intentionally facilitate dialogue across differences and support the professional development of both instructors and students in relation to culturally responsive practices. Students like Charity added to this by emphasizing that all students needed support to unlearn stereotypes and communicate across cultural difference. After describing how she noticed that some White peers tend fake niceness because they don’t want to appear racist, Charity also shared:

> Minority groups [also engage in] generalizing for like… assuming [a White person] is gonna have all these thoughts about you or that they’re going to be racist for instance. ‘Cause it’s more of like judging from person to person and like, not really like placing them in this category as oh, all White people are this or that or just because like maybe their parents might be a certain way doesn’t mean that they are necessarily that way. I feel like for White people they also have challenges in that way knowing that people might already assume these things of them ’cause it works both ways.

Charity highlighted the need for everyone to learn how to engage with each other as individual humans. Everyone holds negative assumptions about one another, and we all need to develop a more empathetic and culturally responsive approach to work together.

While participants did not always explicitly ask to provide professional development in cultural competence for students, this was likely because they were asked to give their *science* instructors advice–– and clearly they would not expect faculty to be able to teach about racism. When considering what advice to provide their instructors, students were very thoughtful and considerate about what would actually be feasible for instructors to implement–– especially those with large class sizes. They shared several strategies, of varying degrees of difficulty, many of which have already been reported in education scholarship. Overall, these findings indicate that there is no need to “reinvent the wheel” as all advice from students have been supported by extant research on effective practices.

## DISCUSSION AND IMPLICATIONS

Overall, the microaggressions described in our findings gives us a clear answer to the first research question. Through counterstorytelling, our participants shed light on how everyday racism manifests in interpersonal interactions in active learning science classrooms at a PWI. Microaggressions stemmed from stereotypes, biases, and assumptions influenced by macro-level structures, such as anti-Blackness, White Supremacy, and deficit thinking. Academic science often claims to be an identity-neutral, meritocratic space for students of all backgrounds (Author, 2026; Cech 2014; Leyva et al., 2022; Morton et al., 2023) which likely exacerbates this tension for Black students in science classrooms. We found that epistemic dismissal was common. This occurs when someone’s knowledge, lived experience, or testimony is unfairly devalued or rejected (Fricker, 2007). Relatedly, the double bind was also prominent in our data, with Black women clearly understanding how their intersectional identities positioned them as intellectually inferior and at risk for such dismissal. Feelings of isolation were exacerbated for Black men and women in the sample who reported multiple instances of blatant avoidance from nearby peers after their instructor told them to turn to a partner. These negative experiences can cause real harm for Black students, as they can lead to racial battle fatigue. Supporting faculty and students in learning how to communicate with one another across differences and discussing the harm of microaggressions are important implications of this work. By creating awareness of racial microaggressions, we can then teach, learn, and have conversations about how to disrupt them (Sue et al., 2009). Future research could include examining peer interactions in active learning science classrooms in different contexts such as Hispanic Serving Institutions (HSI), to nuance the how anti-Blackness might manifest in peer interactions across institutional types and different peer groups.

Black students also shared stories of inclusion and affirmation in peer interactions, centered in community, commiseration, and true collaborative learning. Microaffirmations and counterspaces helped students to feel that their intellectual contributions were taken up and valued, their racialized experiences were recognized and helped them feel protected from racist interactions in small groups. However, the vast majority of microaffirmations reported by participants were only experienced with other people of color, in spaces created for them, and by them, using their community cultural wealth. Therefore, by leveraging their resistant, social, linguistic, navigational, and aspirational capitals, Black students were able to navigate racialized exclusion and built community with peers in their classrooms. While Black students were able to successfully facilitate these interactions using the tools and skills they possess, the onus should not be on them to do so.

Clearly, despite Black students’ strength and resilience, navigating these experiences involve various types of harm–– from the microaggressions themselves, to the rumination and emotional labor to make sense of and navigate these experiences, and trying to create spaces where they would be protected from racist interactions (Solórzano & Peréz Huber, 2020). This has consequences for the student’s cognitive load, affect sense of belonging, and lead to stereotype threat (Steele, 2011; Sue, 2010). Further, the increase of the salience of negative racial/ethnic stereotypes put students of color in a position to have to engage in stereotype management (McGee & Martin, 2011). Regardless of the high achievement they maintain, having to constantly navigate these microaggressions and other racist experiences can lead to racial battle fatigue (McGee, 2021). Therefore, it is important to address this issue in our active learning classrooms. While discussing racialized interactions might feel uncomfortable to us, we need to recognize how much worse it is for students affected by them. Therefore, if bringing attention to these matters can make students feel safety and comfort in their courses, not having to worry about harm or wonder about intent, we should all be willing to reflect and learn about what we can do.

Despite all of this, participants believed active learning was better than lecture because it provides opportunity to co-construct knowledge and address confusions with their peers, share resources and advice, and build community and friendships. Students clearly understood the value of peer discussions in active learning, even as it failed to actualize its full potential in their own peer interactions. Our participants shared various pieces of advice on how to improve active learning for them, naming various practices that have already been shown to be supportive. They did this in part by drawing from positive peer interactions and active learning experiences curated by thoughtful instructors in and out of STEM. Other pieces of advice offered unexpected nuance to issues such as how to best create groups and the need to explicitly teach about group work generally and provide specific support and training around building empathy and cultural competence. We should be co-constructing the classroom with our students, who are the ones experiencing the learning. Implementing suggestions from students will also make them feel a deeper connection with their instructors as this reminds students of their agency in the classroom.

Finally, our findings also point to the importance of formal and informal counterspaces at the university level. In the current political landscape when federal and state governments are releasing “anti-DEI” mandates that effectively dismantle formal counterspaces and affinity groups, it is particularly important that we support the students in our classrooms by intentionally designing our learning environments to center racial equity and intentional cross-cultural community building. One important implication for administrators is to provide faculty and students with culturally competent and responsive trainings. To do this, they can leverage existing interdisciplinary expertise at their institutions such as those in Communication Studies, Cultural Studies, Education, and Sociology, to name a few, and develop/pilot curriculum that addresses these issues at the start of the semester, so that all students are well equipped to engage in supportive, affirming, and equitable peer interactions.

## CONCLUSION

To our knowledge, this is the first study characterizing negative and positive racialized interpersonal experiences of Black students during the peer discussion/group work portions of their active learning college science courses. We found that Black students are being subjected to racial microaggressions that manifested in various ways and can affect their sense of belonging, cognitive load, and ultimately lead to racial battle fatigue. Yet, Black students also leveraged their community cultural wealth to develop counterspaces within their classrooms and make use of extant counterspaces in the institution, to intentionally create affirming learning environments for themselves. Importantly, despite negative experiences, Black students still believed in the value of peer discussion in active learning over traditional lecturing. Finally, participants shared advice for instructors on how to better support racial equity in peer discussion.

We hope to have made clear that the point of this study is not to avoid peer discussions or group work. These participants shared, in no uncertain terms, that peer discussion is more supportive for their learning than traditional lecturing. DBER scholars have also successfully argued that lecturing is detrimental to learning and belonging in and of itself, and even tantamount to discrimination (Hadelsman, 2022). Instead of refraining from using peer interactions in active learning, we must instead acknowledge that there is room for improvement and collectively work towards change. We urge colleagues and institutions to intentionally design active learning environments with racial equity in mind. Not meeting this need will allow racialized harm to persist for students who, like everyone else, are simply trying to learn science and fulfill their academic and personal goals.

## CONFLICT OF INTEREST DISCLOSURE

All authors declare no conflicts of interest.

## ETHICS APPROVAL STATEMENT

This study was approved by University of Georgia IRB #PROJECT00006761

## DATA AVAILABILITY STATEMENT

The data supporting this study are not available to protect participant anonymity and privacy.

## FUNDING STATEMENT

We thank UGA’s Franklin Forward Seed grant for member checking funding.

## Supporting information

Interview Protocol

## ACKNOWLEDGEMENTS

We thank the ACCESS Lab team and BERG for feedback on various iterations of this manuscript.

## Notes

### Competing Interest Statement

The authors have declared no competing interest.

