## Supplementary material for "“Let’s Be Black Excellence”: How Black Students Navigate Exclusionary and Affirming Racialized Peer Interactions in Active Learning": Interview Protocol

***Semi Structured Interview Protocol***

**INTRODUCTION**
Thank you for taking the time to participate in our research project! As a reminder, your participation is voluntary and confidential. Should you feel uncomfortable at any point, you can skip any question or stop the interview at any time. This interview should last up to an hour.

**Before we begin, I want to clarify a few things.**

**First, it is how we are defining active learning in this study. Hand the printout with definition of peer discussion in AL***.**

Do you have any questions about this definition?

**Second, I want to encourage you to pause and think and take your time answering the questions.** We allotted one hour for this interview so that you can do this and not feel rushed.

**Finally, this will feel like a one-sided conversation,** so expect that you will be doing most of the talking. I will be asking questions from a list and ask you to elaborate at times to get more details about your perspectives or experiences. It may get repetitive at times, but it is to ensure we can capture all your thoughts and experiences around these issues.

**Do you have any questions?**

***I'm going to start recording now.***
**START INTERVIEW: Turn on recorder.**

- Now that I've turned on the recording, can you confirm that you give your consent to be recorded?
- To protect your anonymity, when we report our findings, we will use a pseudonym (a fake name). Please choose a fake, human name for us to use as your pseudonym___________________________________

**Warm up questions**

1. In your survey you said you are a ______year, majoring in ______________________.
2. What are your current career goals?
3. In the survey you mentioned the following courses that use active learning. Since I've just clarified what this type of format looks like, do you want to add or remove any of the courses you listed?
   ____________________________________________________________________________
   ____________________________________________________________________________
   ____________________________________________________________________________
   ___________________________________________________

**The purpose of this study is to learn about positive and negative experiences of with students in courses that require peer-peer interactions. We want to learn what works well and what can be improved so that all students can make the most of active learning opportunities.**
**We are going to start with the negative and then move to the positive.**

4. **Please give me some specific examples of negative interactions with classmates in science courses that used active learning.**

*[Potential follow up probes:*
*What happened to make you feel this way?*
*Did someone make you think that?****
*If from self-blame for not participating or not knowing how to interact, probe if they feel that other people in their classes felt the same way.*
*If they mention climate, culture, systemic issues [coming from outside], probe for clarity*
*e.g. Why do think this happened? Can you describe the culture or climate that led to these interactions?*
*If they state they have never had a negative interaction ask: Have you ever witnessed a bad group interaction in your group and if you did, did do you something about it?].*

5. In your demographic survey, you identified as (list all identities): ___________________
_____________________________________________________________________

**How, if at all, do think that one or more of these identities played a role in negative interactions with peers?** Tell me more. *[Make sure they elaborate on things like "assumptions", "stereotypes", "culture" etc.]*

**Tell me more about those peers you had negative interactions with**:

a. Were you working as pair or group?
b. Did you get to pick your group members?
c. How diverse was the group (men, women, race/ethnicity)?

6. Tell me about how you process these negative experiences. What do you do? Tell me more…

7. Are there other students or people whom you trust that you can turn to?

8. When you are interacting with peers in your science courses, are you ever reminded of any of your identities?

9. In your demographic survey, you identified as *(list identities again).* Do you think that people from identities **different** than yours have different negative experiences in peer discussion? How so? Why do you think this is? Tell me more. How about this other group?

10. How, if at all, did the course instructor (professor or TA or PLA) try to support and guide you and your classmates in learning how to work together? (For example, did they assign roles for each group member, give guidelines on how to work together, checked in to see if groups were working well?)

**Thank you for sharing your experiences so far... Now, let's transition to thinking about positive experiences in active learning in your science classrooms.**

11. Please give me some examples of how you have been **encouraged or affirmed or supported** by classmates in science courses that use peer interactions. Tell me more. [*Follow ups:* What happened to make you feel this way? Did someone make you think that? *Probes:**If they mention class climate or culture [coming from outside], probe for clarity—why do you think things were structured like that? How did that culture or climate support these positive interactions?]

12. How, if at all, do feel that one or more of your identities played a role in your positive interaction with peers? Why? Tell me more.

**Tell me more about those peers**:

a. Were you working as pair or group?
b. Did you get to pick your group members?
c. How diverse was the group (men, women, race/ethnicity)?

13. How, if at all, did the course instructor attempt to support and guide you and your peers in learning to work in pairs or groups? (e.g. did they assign roles, give guidelines, have group-check ins?).

14. In your demographic survey, you identified as ______________________________________________. Do you think that people from *identities* different than yours have different positive experiences in these settings? How so? Why do you think this is? Tell me more. How about this other group?

15. In the survey, you mentioned that you attended _____________________High School. How were the peer interactions you described at UGA similar or different than your experiences with peers at your high schools? Make sure to get *demographics of school and whether that informs their experience navigating a PWI.*

16. Given what you've shared today, do you feel that peer discussion in active learning is beneficial to your learning?

17. If you were to give **<u>three pieces</u>** of advice for your course instructors to help improve peer interactions what would those be?

18. Final question: Is there anything you would like to talk about that did not come up, or something you want to circle back to?

**CONCLUSION**

This concludes our interview! Thank you very much for taking the time to share your experiences with me! As the informed consent stated, we also want to conduct a "member check" with you. We would meet with you for about 30 minutes and give you a printout of the transcript of your interview so that you can approve it or add to it. We would provide you with a $20 gift card for your time. Would you be interested in doing this?

*****Study's Definition of "Active Learning"**

We are interested in active learning teaching strategies **where the instructor asks students to pair up or form groups to discuss a topic or work together on a problem set <u>inside class</u>. It is any in-class work that requires that you interact with your peers**. This can be talking with someone sitting next to you to come up with an answer to an impromptu or planned question (TopHat or another poll) during lecture. It can also be a **discussion session** where students work in groups to work over problems on a worksheet. This can include **breakout sessions, discussions, recitations where you are asked to work with peers.**

**Labs <u>do not</u> count as active learning for the purposes of this study**. Neither does group work outside of class. We want you to think about the science courses you've taken here at UGA that used these types of activities inside a classroom.
